# The impact of modern admixture on archaic human ancestry in human populations

**DOI:** 10.1101/2023.01.16.524232

**Authors:** Kelsey E. Witt, Alyssa Funk, Lesly Lopez Fang, Emilia Huerta-Sanchez

## Abstract

Admixture, the genetic merging of parental populations resulting in mixed ancestry, has occurred frequently throughout the course of human history. Numerous admixture events have occurred between human populations across the world, as well as introgression between humans and archaic humans, Neanderthals and Denisovans. One example are genomes from populations in the Americas, as these are often mosaics of different ancestries due to recent admixture events as part of European colonization. In this study, we analyzed admixed populations from the Americas to assess whether the proportion and location of admixed segments due to recent admixture impact an individual’s archaic ancestry. We identified a positive correlation between non-African ancestry and archaic alleles, as well as a slight enrichment of Denisovan alleles in Indigenous American segments relative to European segments in admixed genomes. We also identify several genes as candidates for adaptive introgression, based on archaic alleles present at high frequency in admixed American populations but low frequency in East Asian populations. These results provide insights into how recent admixture events between modern humans redistributed archaic ancestry in admixed genomes.

## Introduction

Admixture, or the genetic merging of two or more parental lineages, has been demonstrated as an important contributor to modern human genetic variation (Rudan 2006; Hellenthal et al. 2014; Martin et al. 2017). Examination of human DNA sequence data from the past and present suggests that admixture has been more frequent than previously thought (Korunes and Goldberg 2021). For example, we now have evidence that archaic and modern humans interbred, and that numerous population replacement and admixture events occurred across Eurasia in the past, resulting in the transmission of genes between previously geographically separated populations (Bustamante and Henn 2010; Green et al. 2010; Reich et al. 2010). Many of the genomes of contemporary individuals in the Americas are the result of various admixture events due to European colonization and the Transatlantic slave trade (Moreno-Estrada et al. 2013; Bryc et al. 2015; Ongaro et al. 2019), as well as continuous gene flow from a number of immigrant populations across Europe, Africa, and Asia (Han et al. 2017).

Many modern humans also show evidence of introgression, or the incorporation of alleles from archaic humans like Neanderthals and Denisovans (Green et al. 2010; Reich et al. 2010). Modern humans encountered both groups as they expanded out of Africa, and high-coverage genome sequencing of a Denisovan (Meyer et al. 2012) and multiple Neanderthals (Prüfer et al. 2014; Prüfer et al. 2017; Mafessoni et al. 2020) has helped characterize the archaic variation that remains in modern human genomes. Neanderthal ancestry is also found in some African populations via admixture with Eurasian populations that harbored archaic ancestry migrating back into Africa (Wall et al. 2013). Studies estimate that non-African populations have 1-4% Neanderthal admixture, with East Asian populations exhibiting more Neanderthal introgression than West Eurasian populations (Wall et al. 2013; Sankararaman et al. 2014), perhaps due to more archaic admixture events with ancestral East Asians, but other plausible scenarios have been proposed (Coll Macià et al. 2021; Witt et al. 2022). Denisovan ancestry, however, shows a more varied distribution: Oceanians have by far the most Denisovan ancestry (up to 5%) (Reich et al. 2011; Vernot et al. 2016), and while South Asians and East Asians also have some Denisovan ancestry, European populations have very little (Sankararaman et al. 2016; Witt et al. 2022). In some cases, these past introgression events likely resulted in adaptive introgression (Bustamante and Henn 2010; Green et al. 2010; Reich et al. 2010) – selection favoring archaic variants to facilitate adaptation to novel environments in modern humans (Racimo et al. 2015). Some candidate regions identified as adaptively introgressed include genes responsible for body fat distribution (Racimo et al. 2018), high-altitude adaptation (Huerta-Sánchez et al. 2014; Yi et al. 2010), skin pigmentation, and innate immunity (Racimo et al. 2017). Many of these adaptive archaic haplotypes are not found in all populations but are continent- or region-specific. We, therefore, observe regional differences in the frequency and distribution of archaic alleles, likely based on the geographic and temporal distance from the initial archaic admixture events, and on the past selective pressures and demographic events that a population was exposed to.

Until recently, the landscape of archaic ancestry in modern humans has primarily been studied in the context of Eurasian populations, and consequently much less is known about how archaic ancestry is distributed in populations in the Americas. Notably, these populations also harbor remnants of archaic ancestry (Sankararaman et al. 2014; Sankararaman et al. 2016; Racimo et al. 2017) as they are the descendants of ancestral populations that interbred with archaic humans in the past. While previous studies show that Indigenous American individuals have a similar or slightly higher proportion of archaic admixture to East Asians (1.37% Neanderthal and 0.05% Denisovan ancestry in the Americas, compared to 1.39% Neanderthal and 0.06% Denisovan ancestry in East Asians (Sankararaman et al. 2016)), less is known about how demography and natural selection have affected archaic variation in these populations. Interestingly, it was recently discovered that Peruvians exhibit the largest number of high frequency derived archaic alleles and the largest number of candidate loci for adaptive introgression (Racimo et al. 2017), further demonstrating the value of examining archaic ancestry in the Americas. One reason why many populations in the Americas are mostly excluded from studies characterizing the effects of archaic ancestry is that many populations are admixed (due to European colonization and the Transatlantic slave trade), and components of ancestry have historically been challenging to identify in admixed individuals. This is especially true of contemporaneous modern American populations, in which admixture has occurred relatively recently (Sankararaman et al. 2008). However, these signals of admixture can be indicative of selection (Tang et al. 2007), past demographic events (Vernot and Akey 2015; Wang et al. 2020), and even disease susceptibility (Rudan 2006; Mautz et al. 2019), demonstrating the value of analyzing admixed individuals.

Furthermore, more studies of admixed populations are needed as human populations are tending to become more admixed due to increased mobility, and we need to investigate how admixture shapes patterns of genetic variation. Recent admixture in the Americas likely impacted the frequency and distribution of archaic variants within a population, and studying these populations can provide more insights into the impact of this recent gene flow on admixture from archaic humans. In this study, we investigate how recent admixture has shaped the distribution of archaic variants in admixed populations in the Americas. We compare the admixed American populations in the 1000 Genomes Phase III data (CLM - Colombians from Medellin, Colombia, MXL - Mexican Ancestry from Los Angeles, PEL - Peruvians from Lima, Peru, and PUR - Puerto Ricans from Puerto Rico) (1000 Genomes Project Consortium et al. 2015) to the high-coverage Neanderthal and Denisovan genomes (Prüfer et al. 2017; Mafessoni et al. 2020). We examine how admixture proportions of European, African and Indigenous American ancestry impact the distribution and amount of archaic variants in these populations. We find that recent admixture increased the heterozygosity and the number of autosomal variants, and that the amount of archaic admixture is proportional to the amount of Indigenous American or European ancestry. Although European and Indigenous American tracts in these admixed genomes have approximately equal proportions of Neanderthal variants, Denisovan variants are found primarily in Indigenous American tracts. An analysis of putatively adaptively introgressed regions with high proportions of archaic ancestry in admixed populations suggests selection for genes related to immunity, metabolism, and brain development. This study demonstrates how recent admixture modulates the distribution of archaic variants in modern admixed populations.

## Results

### Effects of Recent Admixture on Modern Genomes

We began examining admixed populations by comparing them to the unadmixed populations sequenced by the 1000 Genomes Project. By replicating Fig 1b from The 1000 Genomes Project Consortium 2015, we confirm that the number of autosomal variants per genome varies greatly across the 26 populations in the 1000 Genomes Project Phase III dataset (Fig 1A). As expected, African individuals harbor the largest number of variants. Interestingly, individuals from the recently-admixed American populations show a broader range in the total number of variants per genome across a population compared to other populations. This is likely due to the individuals within these populations having varying levels of African and Indigenous American ancestry. To explore whether the number of variants correlates with the amount of African ancestry, we examined modern ancestry tracts (African, European, and Indigenous American) that had been previously identified for the admixed 1000 Genomes individuals as defined by Martin et al (2017) (Table 1, Fig S1), which used RFMix, a random forest-based ancestry inference method to identify haploid ancestry tracts (Maples et al. 2013). African populations and admixed populations with high African ancestry percentages (ACB: African Caribbean in Barbados, ASW: African Ancestry in Southwest US, and to a lesser extent PUR) have the greatest number of variants (Fig 1B). In addition to calculating the number of variants, which counts the total number of sites in the genome containing alternate allele, we also calculated the number of heterozygous sites for each individual. Individuals of recently admixed populations from the Americas also tend to show higher proportions of heterozygous sites compared to South Asian, European or East Asian populations (Fig S2), likely because admixed populations can contain alleles unique to multiple geographic populations (Kidd et al. 2012). This is consistent with theoretical studies that have suggested that admixture between populations can increase heterozygosity in admixed individuals (Boca et al. 2020). Within individuals, the proportion of heterozygous sites in a genome region is also higher in regions containing African ancestry compared to regions with no African ancestry (Fig S3).

**Figure 1.**
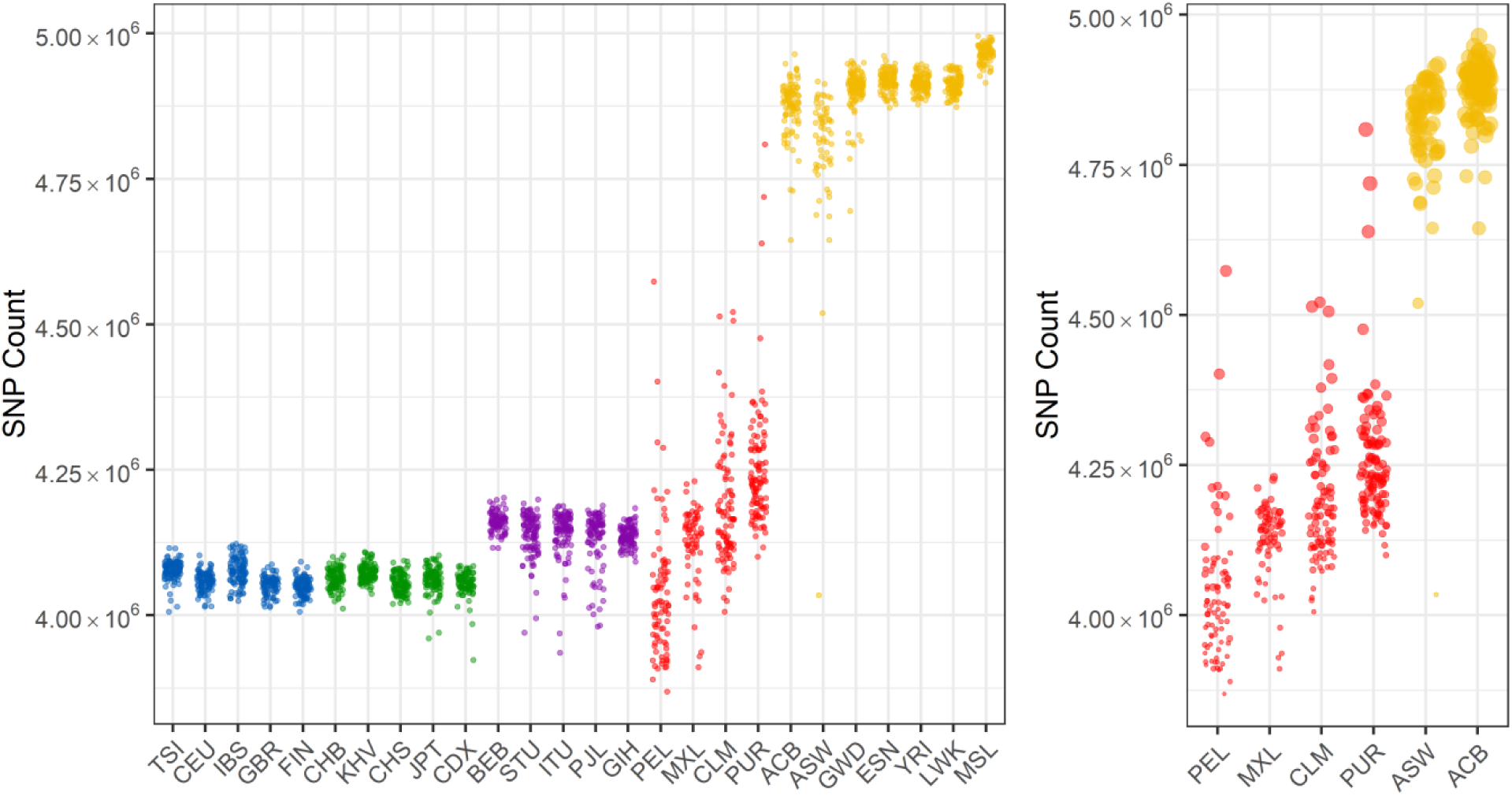
A count of sites containing at least one copy of the alternate allele for each individual. The colors represent superpopulations (blue: European, green: East Asian, purple: Southeast Asian, red: admixed American, yellow: African) a) Counts for each individual in the 1000 Genomes dataset, separated by population. b) Counts for each individual in the admixed American populations, where dot size is proportional to the amount of African ancestry.

**Table 1.**
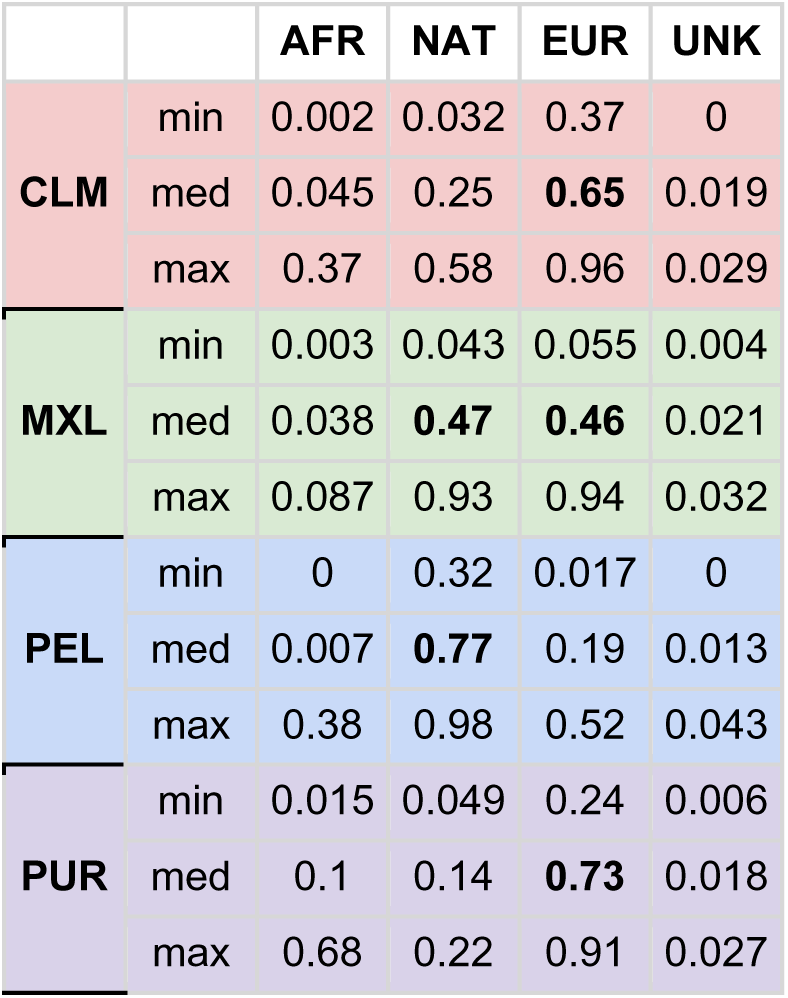
Ancestry in admixed American populations. This table gives the minimum, median, and maximum recent ancestry for each admixed American population analyzed in this study. The values were calculated using the individual data from (Martin et al. 2017). Values in bold show the largest modern ancestry component for the admixed population.

|  |  | AFR | NAT | EUR | UNK |
| --- | --- | --- | --- | --- | --- |
| CLM | min | 0.002 | 0.032 | 0.37 | 0 |
|  | med | 0.045 | 0.25 | **0.65** | 0.019 |
|  | max | 0.37 | 0.58 | 0.96 | 0.029 |
| MXL | min | 0.003 | 0.043 | 0.055 | 0.004 |
|  | med | 0.038 | **0.47** | **0.46** | 0.021 |
|  | max | 0.087 | 0.93 | 0.94 | 0.032 |
| PEL | min | 0 | 0.32 | 0.017 | 0 |
|  | med | 0.007 | **0.77** | 0.19 | 0.013 |
|  | max | 0.38 | 0.98 | 0.52 | 0.043 |
| PUR | min | 0.015 | 0.049 | 0.24 | 0.006 |
|  | med | 0.1 | 0.14 | **0.73** | 0.018 |
|  | max | 0.68 | 0.22 | 0.91 | 0.027 |

### The effects of admixture on archaic variation

Before looking at admixed populations specifically, we examined whether the distribution of archaic alleles varies depending on the geographic location of a population. We used the set of archaic alleles found using sPrime (Browning et al. 2018) and calculated the number of sites with archaic alleles (all archaic, as well as Neanderthal-unique and Denisovan-unique) found in each non-African individual in the 1000 Genomes dataset to examine how archaic site counts differed between populations. We found that East Asians have the greatest number of sites containing archaic alleles, followed by South Asians, while Europeans have the fewest (Fig 2a), as has been previously established (Sankararaman et al. 2014; Sankararaman et al. 2016). The same is true for Neanderthal-unique alleles (Fig 2b). All populations sampled have an order of magnitude fewer Denisovan-unique variants than Neanderthal-unique variants (Fig 2c), and South Asians and East Asians have comparable counts of Denisovan-unique alleles, while Europeans have half that number. Similarly to what we observe for all variants, admixed populations from the Americas have a greater range of sites containing archaic alleles per individual, while populations within Europe, East Asia, and South Asia show more homogeneity. For all types of archaic variants, PUR has the fewest variants, followed by CLM, MXL, and PEL, likely due to differences in the amount of Indigenous American ancestry in these populations: PEL exhibits the highest amount of Indigenous American ancestry (∼75% average) and PUR has the smallest amount (∼14%) (Table 1, Fig S1).

**Figure 2.**
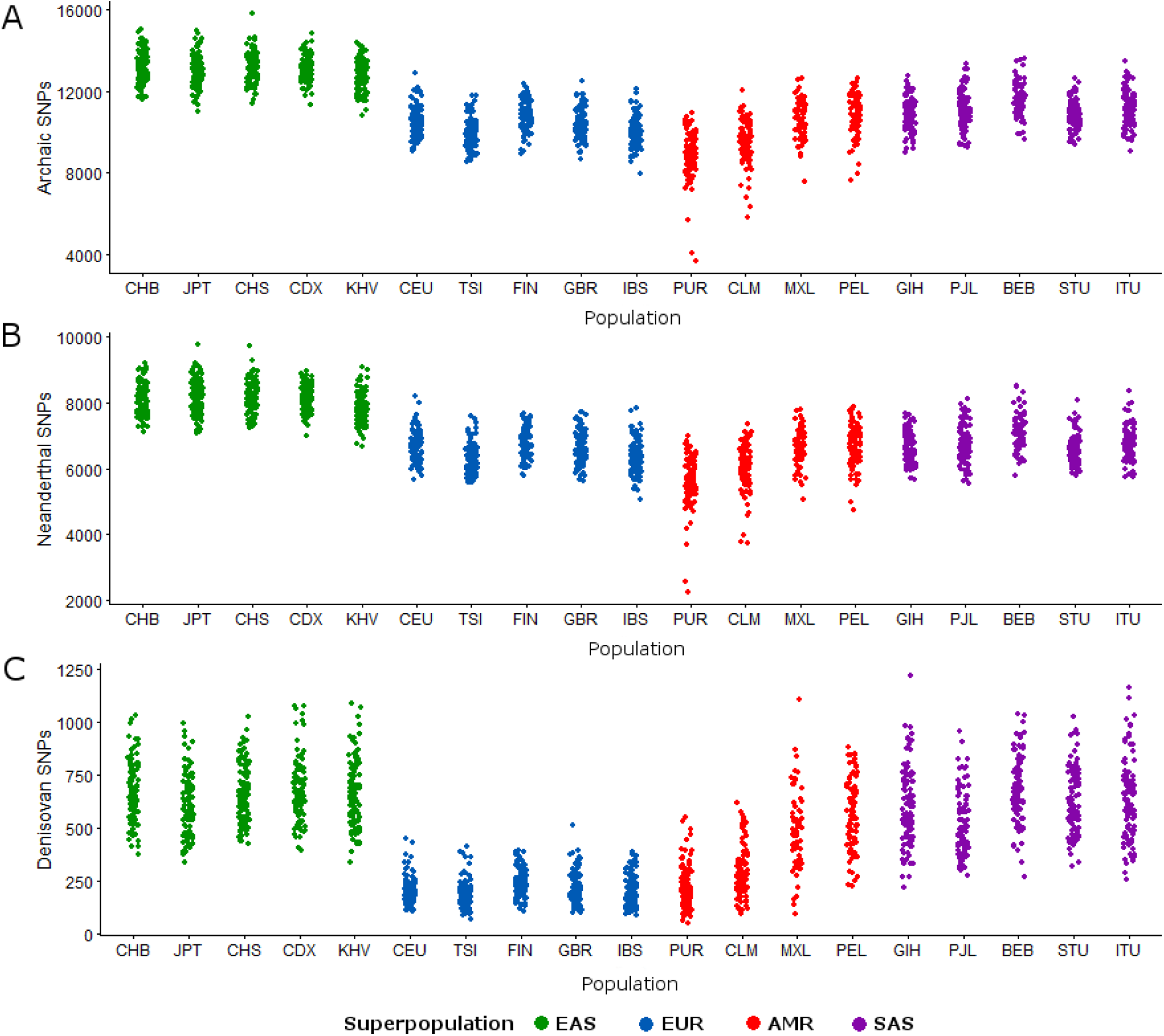
Counts of sites containing archaic alleles (Browning et al. 2018) for all non-African individuals in the 1000 Genomes Project. Color-coding is by super-population, as mentioned in Figure 1. a) Counts of variants found in Neanderthals or Denisovans; b) Counts of variants unique to Neanderthals; c) Counts of variants unique to Denisovans

To explore a possible correlation between the number of Neanderthal and Denisovan alleles and the proportions of modern ancestry in admixed populations, we examined how archaic ancestry was correlated with the amount of African, European, and Indigenous American ancestry in each individual, as defined by Martin et al (Martin et al. 2017) (Table 1, Fig S1). All populations show a positive correlation between Indigenous American ancestry and the total archaic allele count, and a negative correlation between African ancestry and archaic alleles (Fig 3, Table S1). This is expected given that Africans harbor little to no traces of Neanderthal or Denisovan introgression. Interestingly, European ancestry is positively correlated with archaic allele counts in populations with lower amounts of Indigenous American ancestry (PUR and CLM) but is negatively correlated with archaic allele counts in populations with higher proportions of Indigenous American ancestry (PEL and MXL). This is likely due to the fact that Europeans have less archaic ancestry than Indigenous Americans (Sankararaman et al. 2016). For individuals with only a low percentage of Indigenous American ancestry and a high percentage of European ancestry (e.g. PUR and CLM), however, the difference in archaic ancestry proportion between Indigenous American and European ancestry segments has a negligible effect on the total archaic ancestry in each individual.

**Fig 3.**
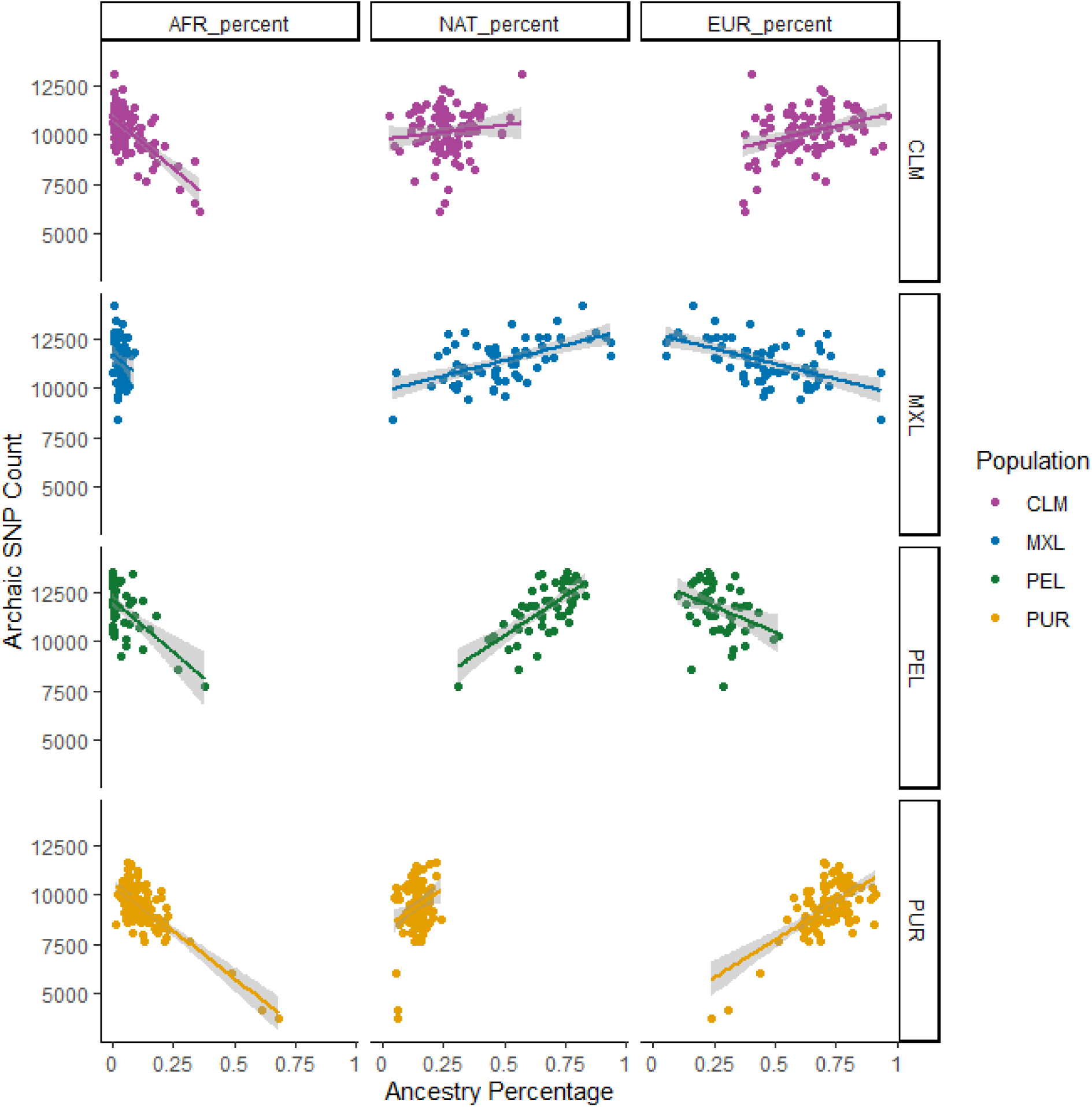
Correlations between percentages of African (AFR), European (EUR), and Indigenous American (NAT) ancestry and count of SNPs containing archaic alleles, subdivided and color-coded by population. The x axis is the genome-wide proportion of a given ancestry type (columns from left to right are African, Indigenous American, and European ancestry) while the y axis is the number of archaic SNPs identified for each individual using Browning et al (Browning et al. 2018). Each row represents a different admixed American population (tp to bottom: Colombians, Mexicans, Peruvians, and Puerto Ricans).

We also assessed whether archaic variants were more likely to be present in specific regions of ancestry across individuals by counting the number of archaic variants in genome regions deriving from the different ancestries (African, European, and Indigenous American, Martin et al. 2017). We calculated *archaic allele density* in these regions by summing the number of positions with archaic alleles identified in all regions with a given ancestry designation (ie. EUR-EUR for both chromosomes having European ancestry or AFR-EUR for one chromosome with African ancestry and one with European ancestry) and dividing it by the combined length of all regions with that ancestry designation. Consistent with the correlations identified in Fig 3, regions of African ancestry are much lower in archaic allele density than regions of European or Indigenous American ancestry (Fig 4), with little to no archaic alleles present in African regions. Variance in archaic allele density is higher in populations with lower proportions of a given type of ancestry (e.g. Indigenous American ancestry in PUR or European ancestry in PEL), where “genome-wide” estimates are likely calculated with only a small number of tracts. Neanderthal-exclusive variants have an average density that is five times higher than that of Denisovans, and Neanderthal allele density is approximately equal between European and Indigenous American segments of the genome. Denisovan allele density, however, is slightly higher in Indigenous American segments than in European segments in populations with higher NAT ancestry (PEL and MXL). This pattern is also observed when looking at individual ancestry tracts, rather than the density across all of an individual’s tracts of a given ancestry designation - of the tracts in the top 1% for highest density (S4 Fig), only one African tract is present (which is similar to a Denisovan haplotype found in some Indigenous American individuals but likely derives from standing ancestral variation (S5 Fig)), and the Indigenous American tracts with high Denisovan allele density outnumber the European tracts with high Denisovan density. Mean archaic allele density for a given ancestry designation (which is the total number of archaic sites in regions with that ancestry divided by the total length of all regions with that ancestry) is approximately equal across all four admixed populations (Fig 4), with the exception of Denisovan allele density, which is higher in PEL and MXL than in CLM or PUR.

**Fig 4.**
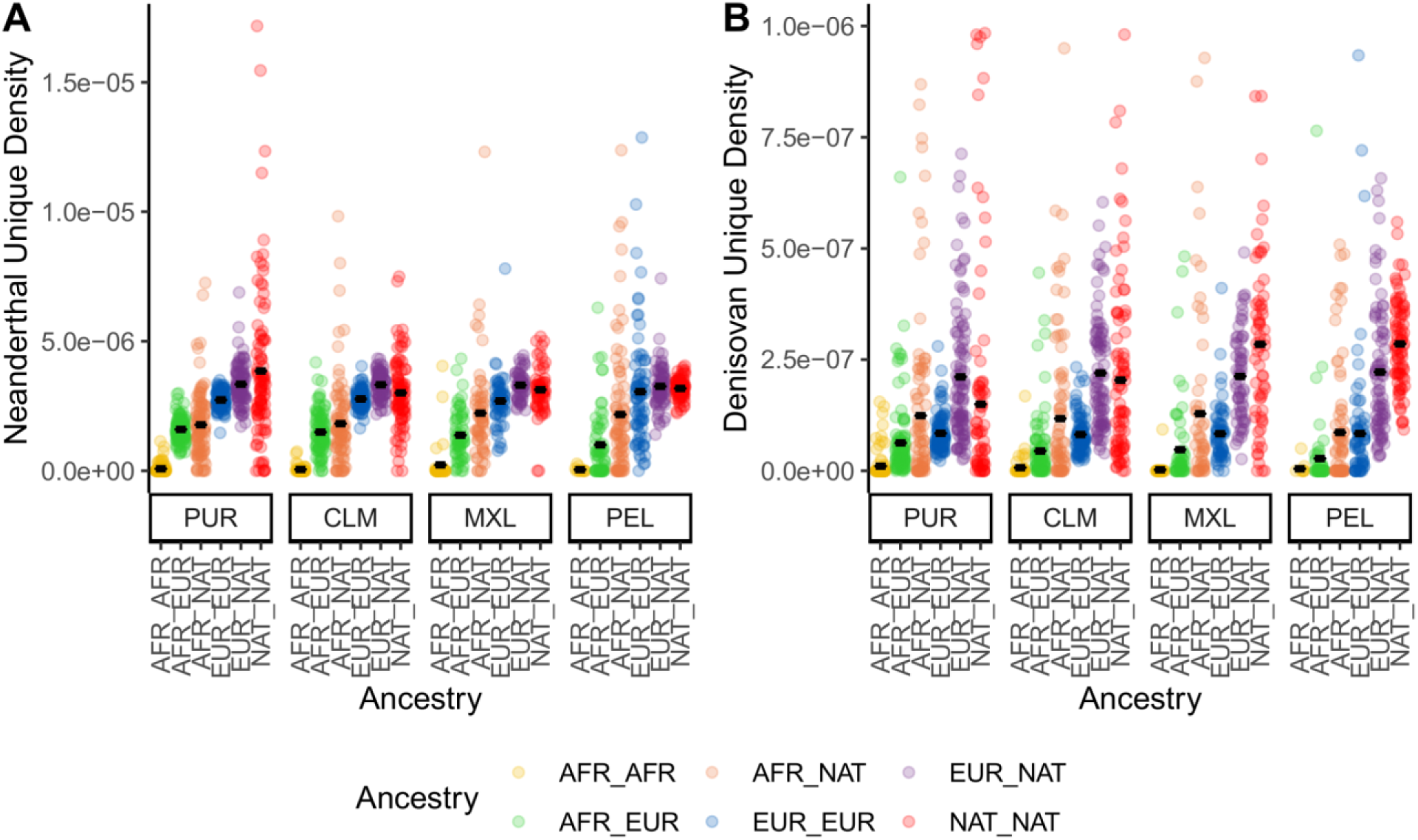
The average archaic allele density across all tracts of a given ancestry type for each individual, subdivided by population. Ancestry abbreviations are ‘AFR’ for African, ‘EUR’ for European, and ‘NAT’ for Indigenous American. Archaic allele density is calculated by subdividing the tracts for each individual into the different diploid ancestry calls (for example, African-African or African-European), summing the total number of positions containing archaic alleles across all tracts of a given ancestry type, and dividing that total by the total length of all ancestry tracts of a given ancestry type. A violin plot shows the distribution, with individual points overlaid on top and a black line to indicate the mean. A) Neanderthal allele density, B) Denisovan allele density

We also examined the Indigenous American ancestry segments specifically for regions that had a high density of archaic alleles and were shared across multiple individuals, to identify candidate regions of archaic introgression that are present at high frequency. We identified a total of five genomic regions that had high archaic allele density in regions of Indigenous American ancestry and were shared between at least 20 admixed individuals: three for Neanderthal alleles and two for Denisovan alleles (Table 2). Of these five regions, two contain smaller segments that have been previously targeted as candidate regions for adaptive introgression (Racimo et al. 2017).

**Table 2.**
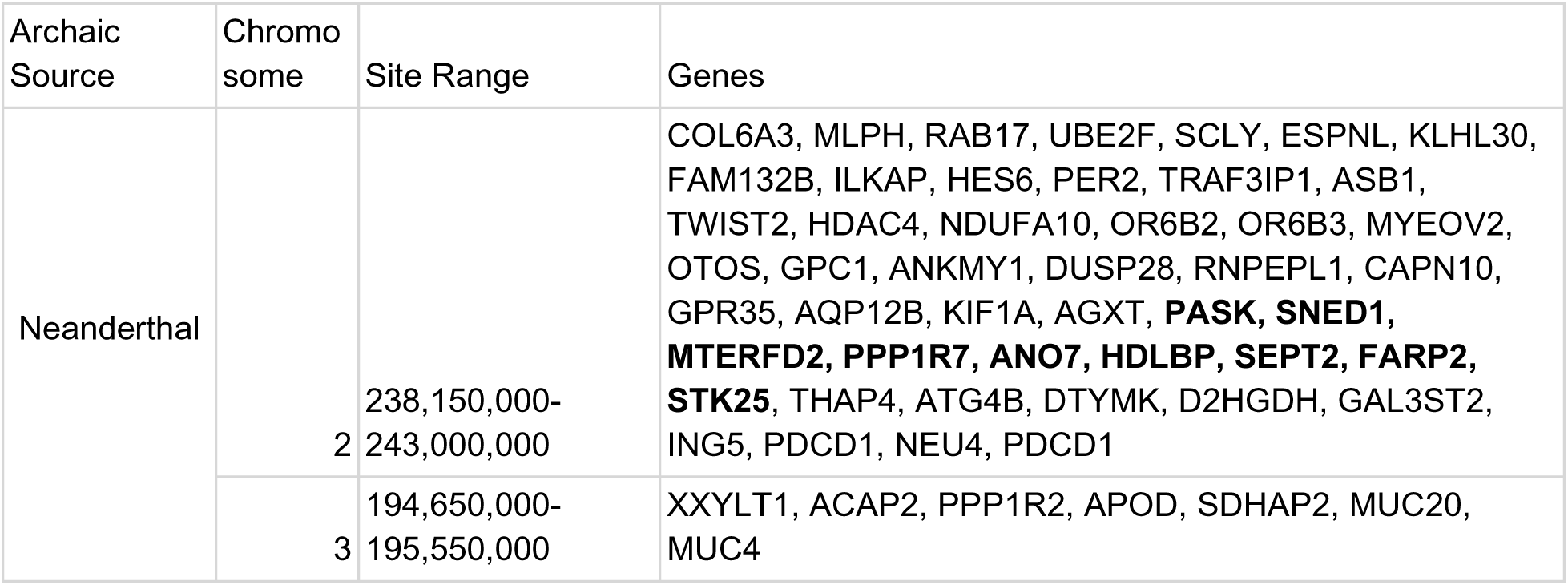
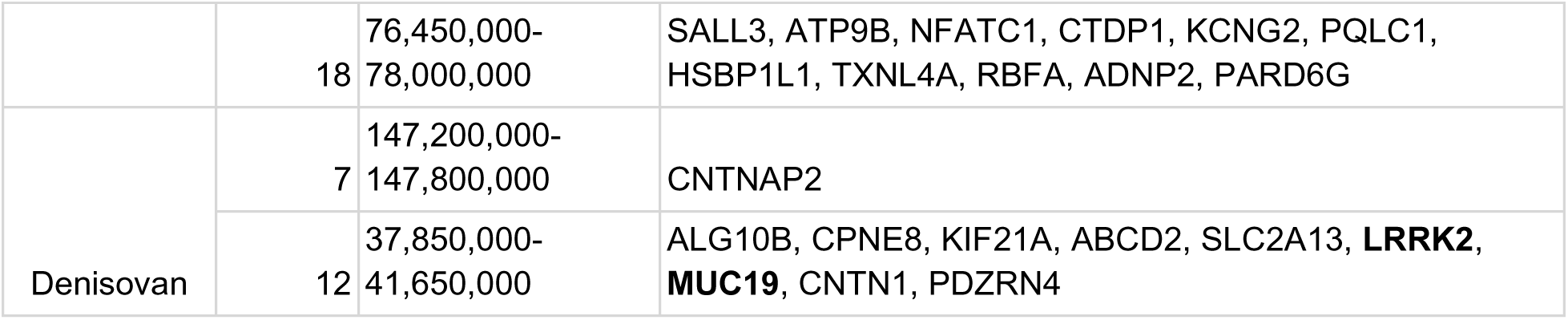
A list of the genome regions with high archaic allele density that are found in at least 20 individuals from PEL,MXL, CLM, and PUR in Indigenous American ancestry tracts. The coordinates correspond to the hg19 genome build. Gene names in **bold** have been implicated as targets for adaptive introgression by other studies (Sankararaman et al. 2016; Racimo et al. 2017). Archaic allele density was calculated for 50 kB windows across the genome, so these genome regions represent multiple consecutive 50 kB windows that had high archaic allele density in Indigenous American ancestry tracts for at least 20 individuals.

### Identifying local adaptation with archaic alleles

We wanted to assess the possibility of adaptively archaically introgressed regions that were locally adaptive to environments in the Americas. If there are loci adaptive specifically for American populations, we would expect them to show signs of positive selection and have archaic allele frequencies in American populations that are higher than the allele frequencies in closely-related populations outside of the Americas. Previous work by Racimo et al. (Racimo et al. 2017) identified a number of 40-kb putative introgressed regions in the 1000 Genomes populations. These candidate regions were identified as having the top 0.1% of number (and frequency) of shared alleles between the test and archaic population that are absent in African populations, and PEL had the largest number of candidate regions identified (Racimo et al. 2017). We chose to use the population branch statistic (PBS) to scan these regions for archaic alleles that show a signal of positive selection. This method requires the use of three populations: the population of interest, a closely-related population for comparison, and an outgroup population. We used PEL and MXL as our Indigenous American populations, as those two populations have the highest average proportion of Indigenous American ancestry of all the admixed populations (Table 1), and used CEU as our outgroup population. Although modern-day Siberians are the closest relatives to modern Indigenous Americans (Malyarchuk et al. 2011; Sikora et al. 2019), we use an East Asian population (CHB) as a proxy for the ancestral population to the founding American populations, which diverged from the Siberian population ancestral to Indigenous Americans approximately 30,000 years before present (Moreno-Mayar et al. 2018; Sikora et al. 2019). We also calculated the archaic allele frequency difference between the American population and CHB to ensure that sites that appear to be positively selected actually have a higher archaic allele frequency in the American population. For comparison, we additionally calculated the allele frequency difference for these populations for alleles that are rare in Africa (f<0.01) but not shared with archaic genomes, to assess whether archaic and non-archaic alleles as a whole have experienced different selective pressures (S6 Fig). New variants that have arisen in populations since humans expanded out of Africa are due either to novel mutation or gene flow from archaic humans, and so we would expect both of these variants to be impacted by the same demographic events, such as bottlenecks. A lower allele frequency difference in archaic variants would suggest that the archaic variants had a more negative impact on fitness.

We were able to identify a number of archaic alleles that have a significantly higher allele frequency in the American populations compared to CHB, which suggests they may have been positively selected (Fig 5). Some of these regions contain genes that have already been identified as adaptively introgressed in Indigenous American populations, including *IFIH1/FAP* (Ávila-Arcos et al. 2020) and *WARS2* (Racimo et al. 2018), identified in PEL, and *LRRK2/MUC19* (Reynolds et al. 2019), identified in MXL. Other genes that were identified as candidates for adaptive introgression include a region identified in PEL and MXL that contains *FARP2* (regulates cytoskeletal formation), a region in PEL and MXL that contains *PAX3* (a transcription factor important to development), a region in PEL and MXL that contains *CNTNAP2* (which affects cell receptors in the nervous system and has been implicated in neurodevelopmental disorders) and a region in PEL that contains *MYOCD*, a gene involved in cardiac function (Fig 6). The distributions of allele frequency differences between American populations and CHB are fairly similar across archaic and non-archaic alleles, and they have similar mean and standard deviation values (S6 Fig).

**Fig 5:**
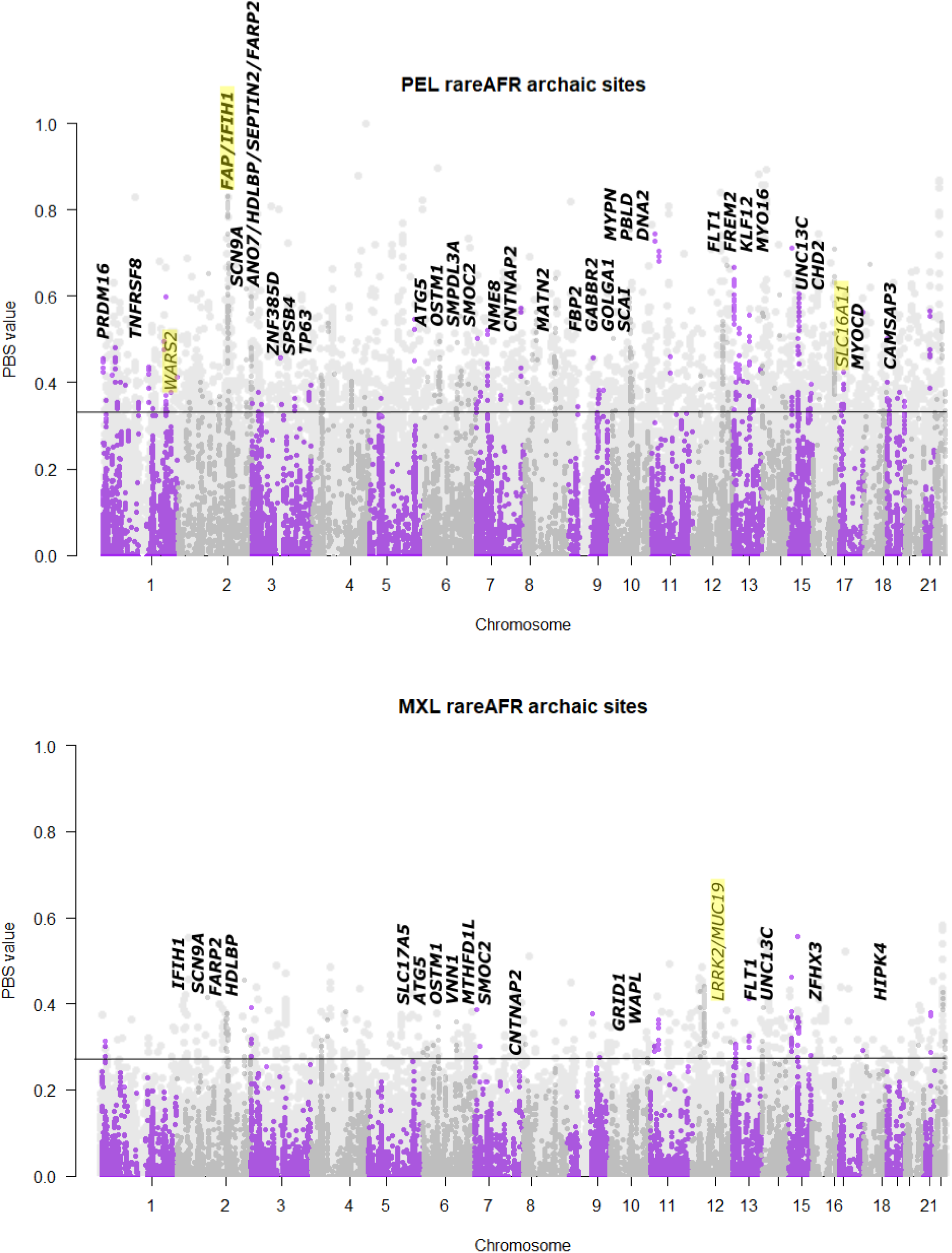
PBS results comparing the American population (A: PEL, B: MXL) with East Asians (with Europeans as outgroup). Bold labels are genes with at least 10 archaic alleles within a gene that have PBS values above the 95th percentile of randomly-selected sites that are rare in African populations and greater than 1% frequency in non-African populations.Italicized (non-bold) gene names have 10 archaic alleles within 100kb of a gene but not in the gene itself. Genes highlighted in yellow have been previously identified as putatively under selection. Light grey background dots show PBS values calculated for all segregating sites across the population’s genome, and the horizontal line shows the threshold for the top 5% of all PBS values calculated at each variant site in genomes for the population. Darker grey and purple alternate to show chromosomes.

**Fig 6:**
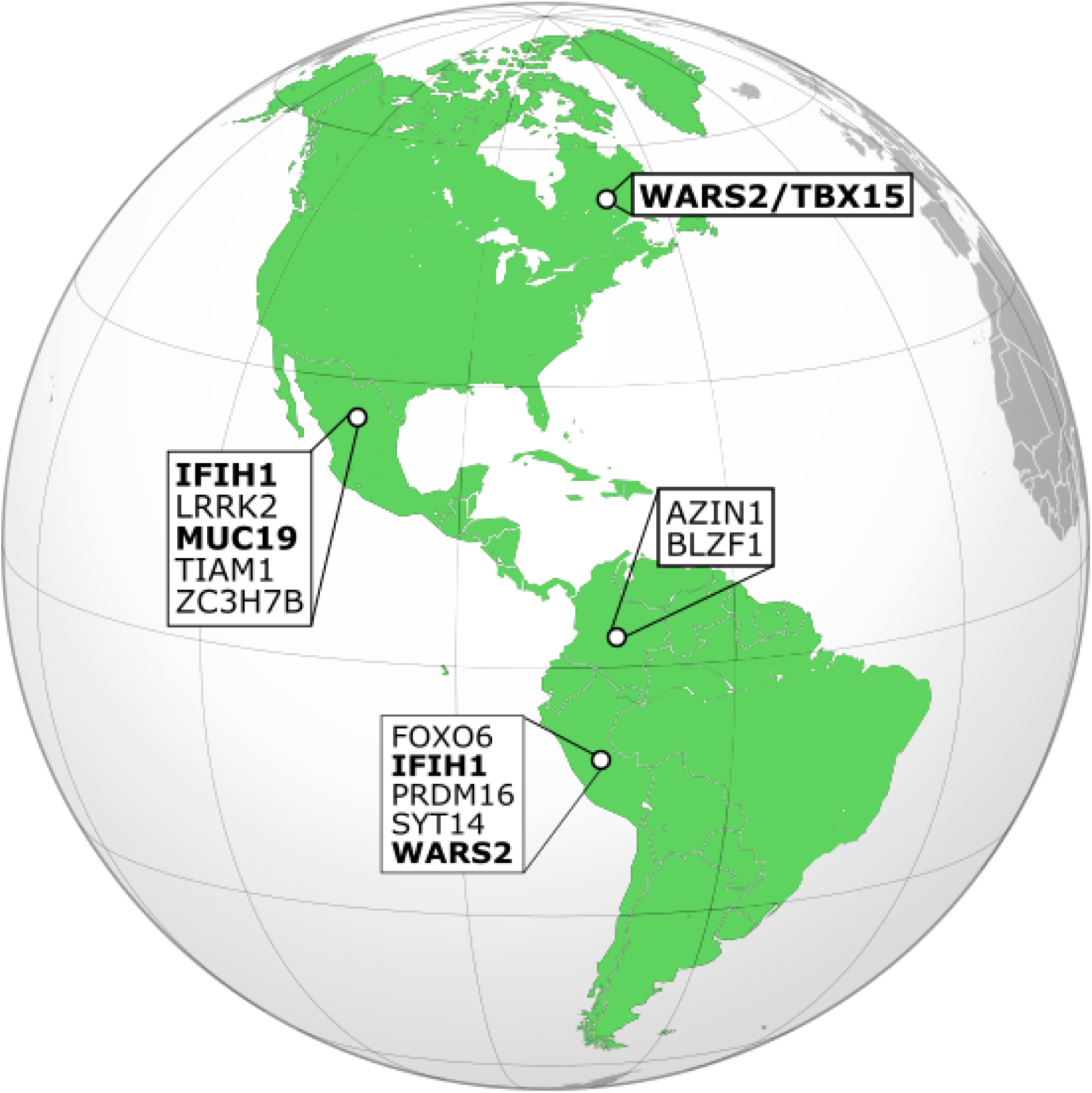
A map depicting the genes identified as adaptively introgressed in American populations. Any genes highlighted in bold have also been identified in other studies.

## Discussion

The increased variant count in African populations is consistent with the out-of-Africa model of human migration across the world, as well as the decreased genetic diversity in non-African populations as a result of the bottlenecks that occurred as humans moved into Eurasia, Oceania, and the Americas (Ramachandran et al. 2005; DeGiorgio et al. 2009; Li and Durbin 2011). Admixed individuals in the Americas show the second highest level of autosomal variants, consistent with their proportions of African ancestry. The admixed populations also show higher variance in the number of segregating sites per individual (Fig 1, Fig 2), as each individual in the population has varying contributions of African, European, and American ancestries.

The amount of archaic variation present in an admixed individual is also impacted by their proportions of African, European, and Indigenous American ancestry. As expected, African ancestry is negatively correlated with the number of archaic variants, and regions of the genome identified as African have few archaic variants, while non-African ancestry is positively correlated with archaic variation. The increased Denisovan variation in Indigenous American ancestry tracts compared to European ancestry tracts suggests that Indigenous Americans have similar archaic allele frequencies to other Asian populations, consistent with their shared history (Dulik et al. 2012; Raghavan et al. 2014). Two populations, PEL and MXL, show a slight negative correlation between European ancestry and Neanderthal variants, suggesting that Indigenous American tracts are contributing more to archaic ancestry than European tracts in populations with high Indigenous American ancestry.

Our analysis of the distribution of allele frequency differences between admixed American populations and East Asians showed that archaic alleles had a comparable allele frequency difference compared to alleles that were rare in Africa but not archaic in origin (S6 Fig). However, a small proportion of SNPs with archaic alleles were at much higher frequency in American populations than Han Chinese individuals, suggesting that positive selection may also have occurred for some alleles. Many of these alleles were over 2 standard deviations from the mean of the allele frequency differences. We used PBS to identify possible candidates for adaptive introgression that were unique to admixed American populations, and identified alleles in multiple genes that have already been identified as targets of adaptive introgression (Racimo et al. 2017; Racimo et al. 2018; Ávila-Arcos et al. 2020), as well as in novel candidate genes, including some that govern development and cardiac function. Some of these genes, including *FARP2*, *MUC19*, and *CNTNAP2*, were also found in Indigenous American ancestry regions with a high proportion of archaic variants (Table 1). The fact that these alleles are found at a high frequency in admixed American populations and that they are specifically found in Indigenous American ancestry segments supports the hypothesis that archaic alleles of these genes may have been adaptive specifically for Indigenous American populations.

The pattern of multiple alleles found at high frequency in a given genomic region that we observed can also be produced by heterosis, where recessive deleterious variants present in both a donor and a recipient population are masked as a result of admixture. When admixture occurs, the higher heterozygosity can mask the deleterious variation, showing a similar pattern to positive selection, especially in regions with high exon density and low recombination rate (Zhang et al. 2020). As heterosis is most likely to occur when recombination rates are low and the exon density in the region is high, we examined the recombination rate and exon density of the possible targets of adaptive introgression (i.e. candidate regions exhibiting high frequency archaic alleles). We found that only two of the SNPs (∼2.4%) were in genomic regions above the 90th percentile for high exon density and low recombination rate (recombination rates of 8.22 x 10^-9^ for *CYP2B6* and 9.16x10^-9^ for *SPRR2F*). This suggests that, given the features of the genome region they are in, the majority of these candidate regions cannot be explained by heterosis.

This study of archaic ancestry in admixed populations illustrates how a recent history of admixture can impact variation introduced from archaic humans. First, by partitioning the admixed genomes into regions corresponding to different ancestries, we find that each modern ancestry component within an admixed population mirrors what we see in the unadmixed ancestral population. For example, archaic ancestry is found almost exclusively in non-African populations, and is nearly absent from genomic regions with African ancestry in admixed individuals as well. This suggests that admixed individuals can be informative about the amount of archaic ancestry in multiple ancestral populations.

However, while we were able to observe the archaic ancestry present in the separate ancestry components of an admixed individual’s genome, they are imperfect representations of the populations the ancestry sources represent. For example, the Indigenous American ancestry segments found in modern individuals likely represent only a fraction of the genetic diversity found in Indigenous Americans prior to European colonization. We would therefore expect pre-colonization Indigenous American populations to have even more archaic variants, especially since Indigenous American ancestry is positively correlated with archaic ancestry in admixed American populations (Figure 3). and by examining the archaic ancestry present in ancient Indigenous Americans we can perhaps identify additional introgressed regions that were targets of positive selection prior to European colonization. We also acknowledge that in this study we focus on populations that all have the same admixture sources and timing, and other admixed populations with a different demographic history (such as ancestry sources that are more closely-related) may not show the same clear differences in archaic ancestry distribution across different modern human ancestry tracts. Future work that examines the changes in archaic ancestry through time in admixed populations, or work that examines populations other than admixed Americans, will help clarify how recent admixture in modern humans affects the amount and distribution of archaic ancestry in the genome.

In summary, we have shown how recent admixture events impacted archaic ancestry in admixed individuals. We find that the proportion of African to non-African ancestry is proportional to heterozygosity and the number of variants, and inversely proportional to the amount of archaic ancestry. We also identify a number of candidate loci that may have been adaptively introgressed, and further exploration of these variants will contribute to a better understanding of the evolutionary processes (such as the timing of admixture and strength of selection) (Corbett-Detig and Nielsen 2017; Zhang et al. 2021) and whether these variants contribute to complex traits in modern humans. Many individuals living today have ancestries deriving from multiple populations, and additional studies to characterize the impacts of admixture on an individual’s genetic variation is needed to gain a better understanding of modern human diversity and its role in disease risk.

## Methods

### Characterizing genetic diversity in 1000 Genomes populations

In this study, we focus on bi-allelic sites at single nucleotide polymorphisms, so any reference to an “allele” would refer to the variant at a single nucleotide position. We aimed to better understand patterns of genetic variation present in the 1000 Genomes Project Phase III dataset by counting the number of autosomal biallelic variant sites for each individual across the 26 populations in the study. To investigate the impact of admixture, we also calculated the African ancestry proportion for each individual in the 6 admixed populations (ACB, ASW, CLM, MXL, PEL, and PUR). The proportion of African ancestry was calculated by summing the lengths of all African ancestry tracts for each individual from the analysis by Martin et al. (2017). If there was an African tract on both chromosomes, then the total length of this tract was used, while if there was an African tract on only one chromosome then half of the tract’s length was used. Lastly, we divided the length of all African ancestry tracts by the total length of all ancestry tracts to account for the small percentage of tracts that had unknown ancestry. For the rest of our analyses, we considered ACB and ASW as African populations due to their high proportion of African ancestry, and so we will use the term “admixed American populations” to refer to the other four admixed populations in the 1000 Genomes dataset (CLM, MXL, PEL, and PUR).

We also sought to uncover levels of heterozygosity, and counted the number of biallelic heterozygous sites for each individual across the autosomes and the X chromosome. When counting on the X chromosome we only considered positions within the pseudoautosomal regions, where males and females are both diploid. To examine the distribution of heterozygous sites within admixed genomes, we divided the genome of the individuals from admixed American populations into regions for each combination of ancestry calls (both chromosomes African, one African and one European chromosome, one African and one Indigenous American chromosome, both chromosomes European, one European and one Indigenous American chromosome, and both chromosomes Indigenous American), according to the ancestry tracts determined by Martin et al. (2017). The proportion of heterozygotes in each ancestry tract for each individual was calculated, and then averaged across all tracts of a given ancestry type for each individual. A Dunn test was used to determine if the distribution of average heterozygote proportions differed for each ancestry type, and a Bonferroni correction was used.

### Characterizing archaic introgression in 1000 Genomes populations

To quantify the amount of archaic ancestry present in each individual and population, we used two methods that employed the counting of alleles that are shared with archaic humans. First, we used the list of archaic sites identified by Browning et al (2018), filtered out all sites that were not bi-allelic, and counted all positions that contained an archaic allele (any position with a “match” to Neanderthal or a Denisovan), as well as Neanderthal-unique (a “match” to Neanderthal and a “mismatch” to Denisovan) and Denisovan-unique sites (a “mismatch” to Neanderthal and a “match” to Denisovan). Second, we compared the 1000 Genomes population data to the Denisovan and Altai, Chagyrskaya and Vindija Neanderthal genomes. The archaic SNP calls were filtered for a minimum genotype score of 40. An allele was then considered “archaic” if it was shared with an archaic individual (Denisovan-unique, Neanderthal-unique, and all archaic alleles were considered) and found at a low frequency in Africa (< 0.01), but had a frequency of 0.01 or greater in at least one non-African population (Witt et al. 2022). We will refer to this method of archaic allele counting as the “allele frequency counting method”. For both methods, allele counts were made per individual per allele (how many archaic alleles were present), as well as per position (how many positions contained archaic alleles).

### Assessing impact of ancestry on archaic allele distribution

To assess whether admixed ancestry impacts the amount of archaic ancestry in an individual, we used the ancestry designations discussed previously (from Martin et al. 2017), and separated the tracts based on the diploid ancestry call (e.g. two chromosomes with African ancestry, one chromosome with African ancestry and one chromosome with European ancestry, etc.). We counted the number of positions containing archaic alleles using Sprime (Browning et al. 2018), including all archaic alleles, Neanderthal-unique alleles and Denisovan-unique alleles in each type of ancestry tract, and calculated the allele density for each ancestry tract in each individual by dividing the archaic position count by the tract length. To calculate the average archaic density for each ancestry type in each individual, we used the following equation:

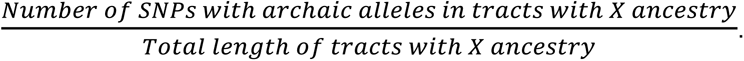

We also compared archaic allele density of individual tracts by examining the ancestry tracts with the top 1% for archaic, Neanderthal, and Denisovan allele density. For this analysis, we focused specifically on regions where both chromosomes had the same ancestry (African, European, or Indigenous American). One tract with African ancestry was in the top 1% for Denisovan allele density (chromosome 10, 10:13980030-15983537, with 9 Denisovan alleles between 14080000-14090000), and we further examined that segment to determine if the ancestry was mis-called or if it was ancestral standing variation. We used haplostrips (Marnetto and Huerta-Sánchez 2017) to compare the Denisovan haplotype to that of American and African populations, and looked specifically at the individuals with 15 or fewer differences from the Denisovan haplotype (S5 Fig).

We determined if Indigenous American ancestry tracts with high archaic allele density were shared between individuals by taking the ancestry tracts with the top 1% allele densities and examining them for overlap of at least 50 kB. If multiple 50 kB tracts all had high allele density, we merged them into one larger tract. The overlapping tracts with the top 5% of greatest sharing between individuals are reported here. This analysis was repeated for all archaic variants as well as Denisovan- and Neanderthal-unique variants - the analysis examining all archaic variants returned the same results as the one analyzing Neanderthal variants, so these regions are reported as being specifically Neanderthal in origin.

To find candidate genes for archaic introgression in each admixed American population, we identified alleles using the “allele frequency counting method” and calculated the population branch statistic between admixed American populations, CHB (the comparison population), and CEU (the outgroup population). We calculated PBS only for sites that had an archaic allele frequency greater than 20% in the admixed population. To calculate F_ST_, we used the Weir and Cockerham estimator (from 1984) implemented by vcftools. PBS for each archaic site was calculated according to the following formula:

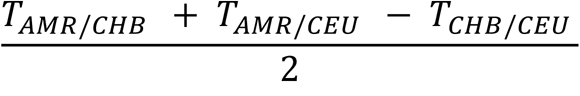

where *T* = –*log*(1 – *F_ST_*) (Yi et al. 2020)

To determine the threshold for outlier archaic sites, we calculated PBS for 2 million random sites across the genome (without replacement) and set the threshold at the cutoff for the top 1% of PBS values for the random sites.

### Identifying regions of local adaptation with archaic alleles

To investigate sites that might contribute to adaptive archaic introgression of genes, we first identified sites with archaic alleles using the “allele frequency counting method” noted above. We used this method to expand the set of SNPs identified as archaic using sPrime, as other studies have indicated that the sPrime algorithm misses some alleles that have previously been identified as archaic in origin (Browning et al. 2018; Zhang et al. 2021). Once those alleles were identified, we calculated PBS for each archaic site using PEL or MXL as the AMR population, CHB as the comparison population, and CEU as the outgroup. We also calculated the allele frequency difference of the AMR population minus CHB for each archaic allele. To calculate a cutoff value for significance, we also calculated PBS and the AMR minus CHB allele frequency difference for all alleles that are rare in Africa but not shared with archaic humans. To determine candidates for local adaptive archaic introgression, we used genes with greater than 10 sites above the mean of the non-archaic alleles plus two standard deviations. We also compared the allele frequency difference between PEL/MXL and CHB for archaic alleles to the allele frequency difference for alleles that were rare in Africa.

## Acknowledgments

KEW and AF are supported by NIH grant to EHS 1R35GM128946-01. EHS is also supported by the Alfred P. Sloan Award.

---

**Fig S1:**
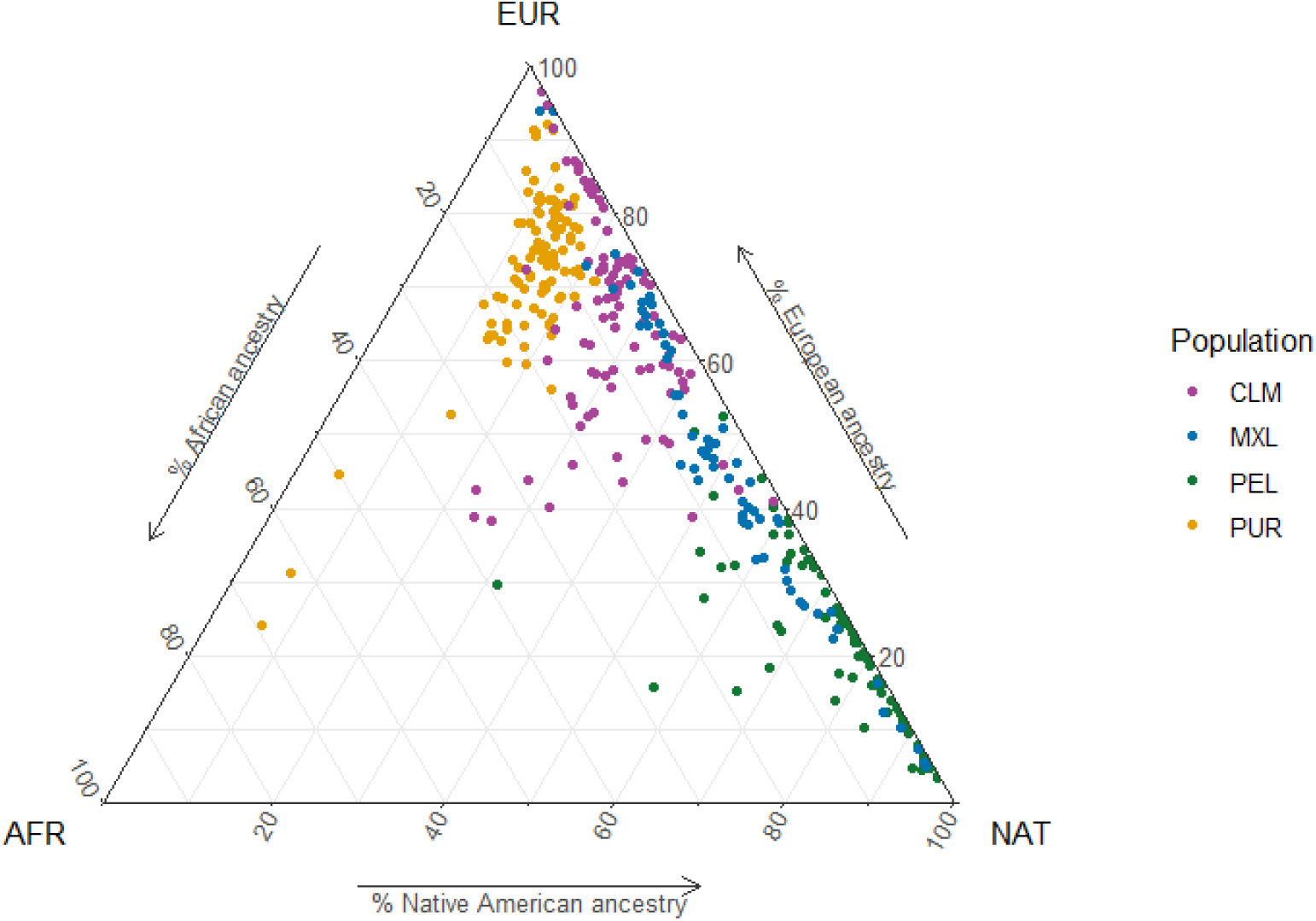
A schematic that illustrates the proportion of African (AFR), European (EUR), and Indigenous American (NAT) ancestry for each of the individuals in the CLM, MXL, PEL, and PUR populations from the 1000 Genomes dataset. Individuals are color-coded by population. Each side of the triangle represents a different type of ancestry, and individual ancestry values are calculated from tracts inferred by Martin et al. (2017)

**Fig S2:**
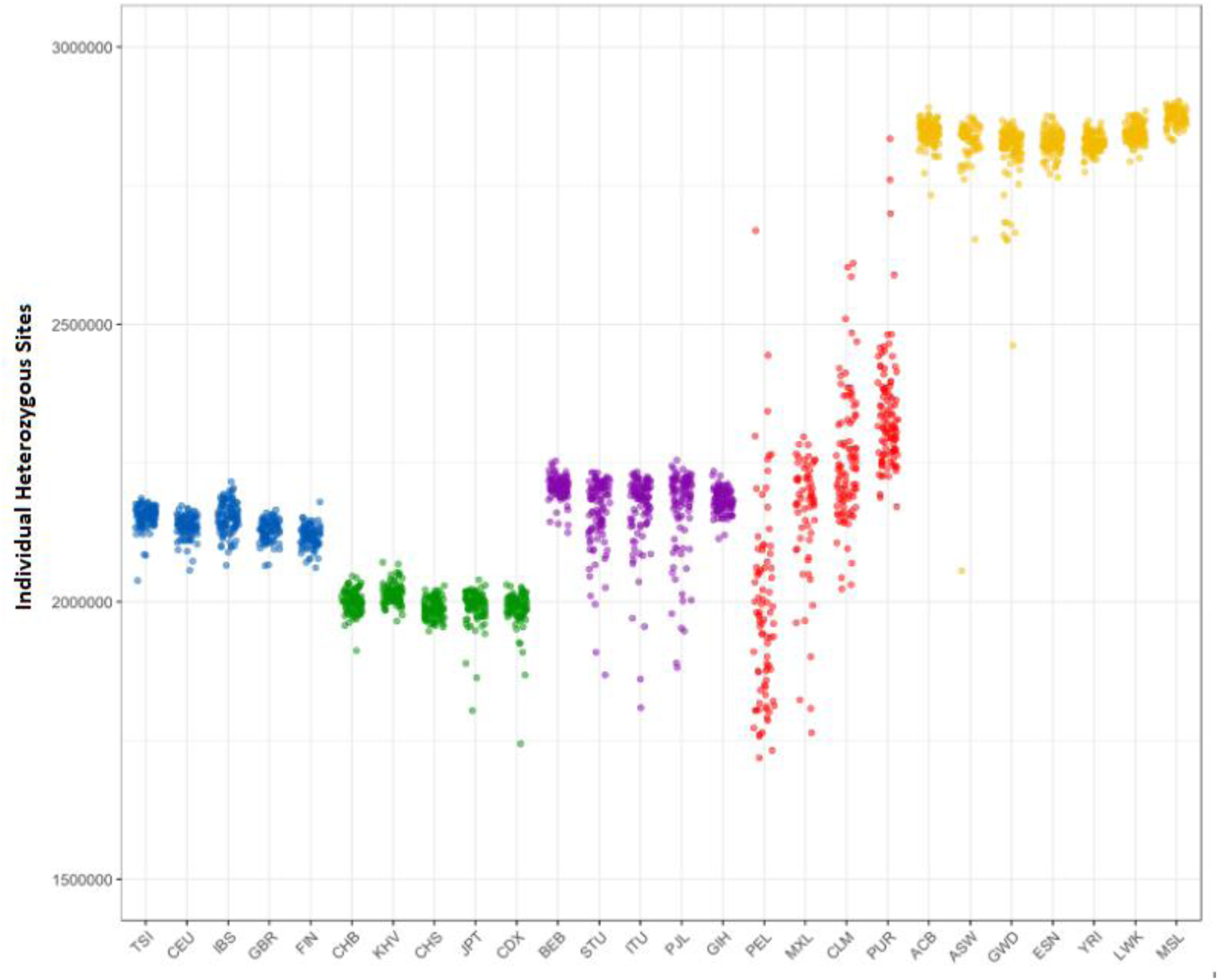
The number of heterozygous sites identified for each of the individuals in the 1000 Genomes Project. The individuals are color-coded by super-population.

**Fig S3:**
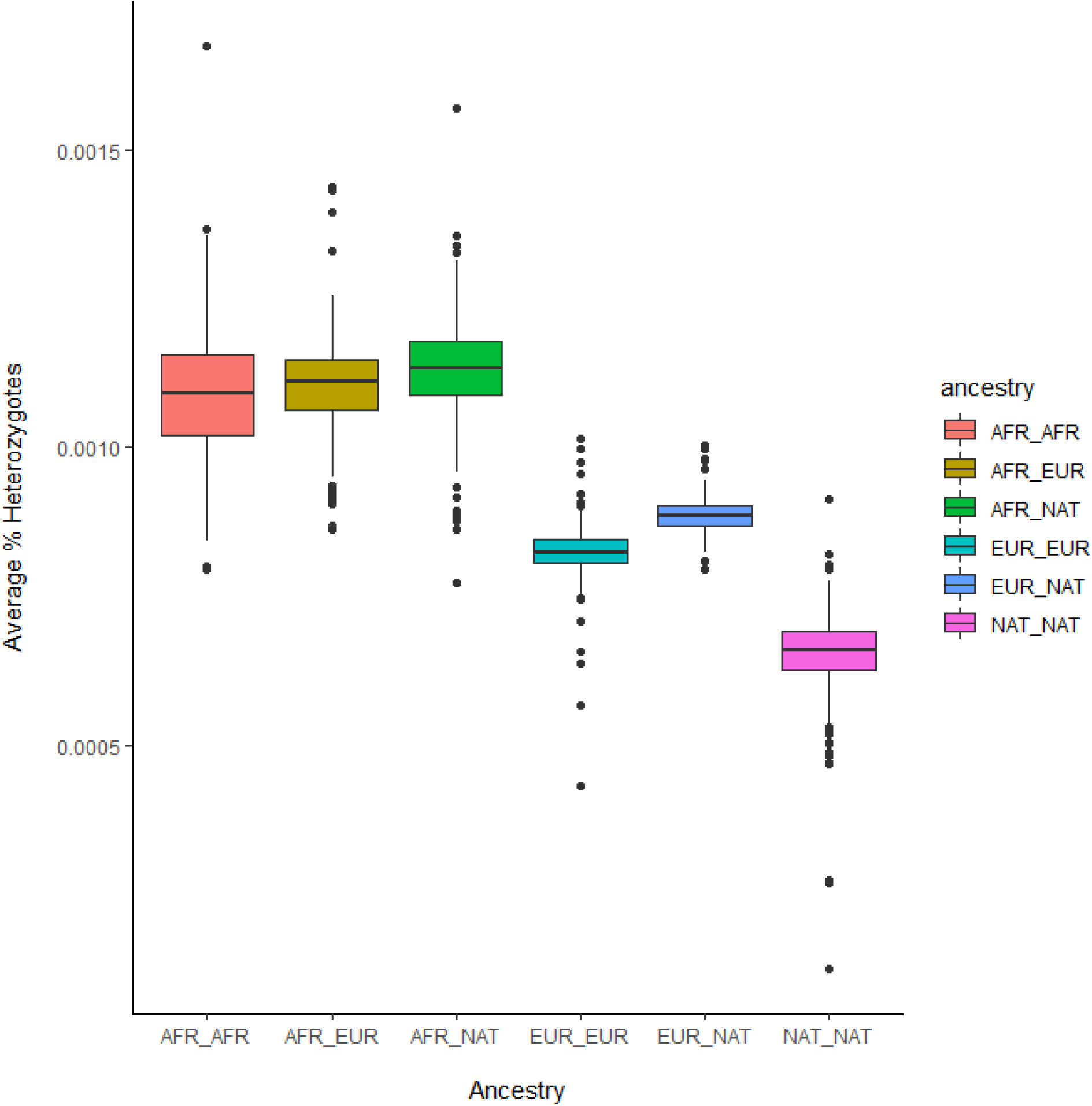
The proportion of heterozygous sites for each diploid ancestry designation for each individual. The average count of heterozygotes was calculated by summing the number of heterozygous sites across every tract with a given ancestry designation, and dividing them by the total length of the ancestry tracts.

**Fig S4:**
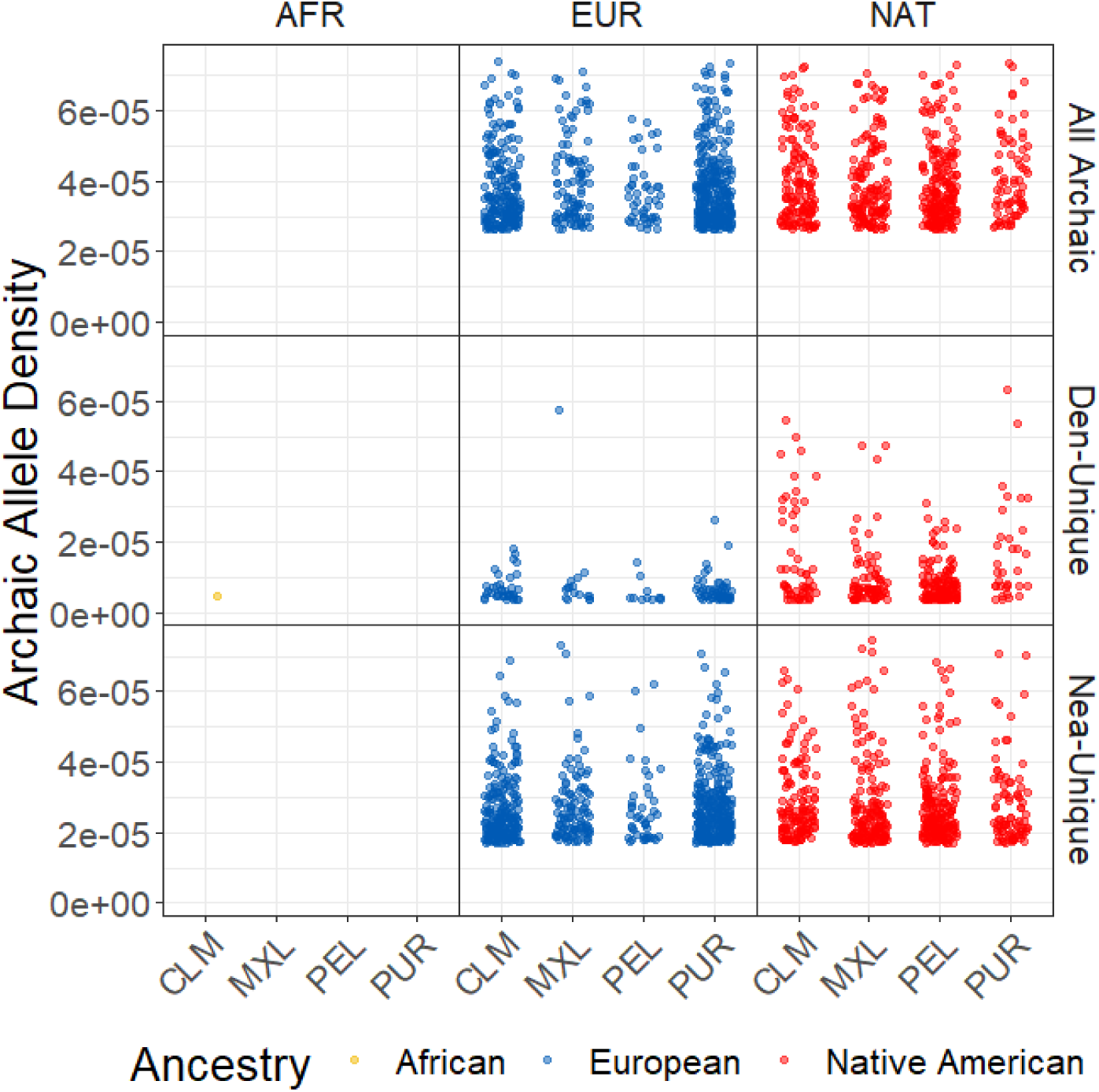
The archaic allele densities of the top 1% of ancestry tracts across all individuals and populations in the Americas. Only regions with homozygous ancestry calls (such as European-European) were considered for this analysis. From top to bottom, the plots reflect Neanderthal-unique allele density, Denisovan-unique allele density, and all archaic allele density. All of the top 1% of tracts had European or Indigenous American ancestry with the exception of one African segment, which had a high proportion of Denisovan-unique ancestry.

**Fig S5:**
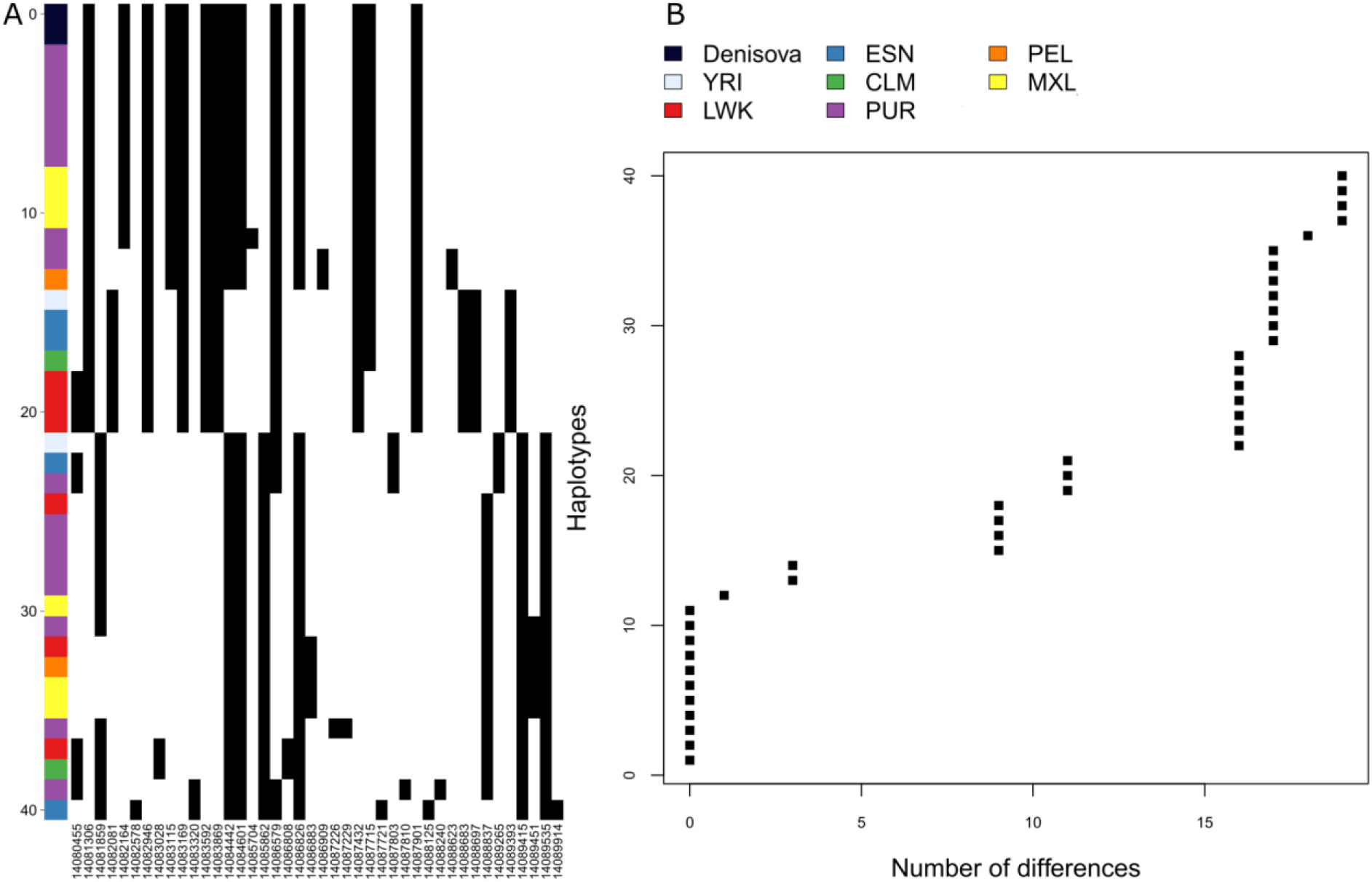
A Haplostrips plot of the African and American haplotypes most similar to the Denisovan haplotype for the African ancestry tract that was in the top 1% for Denisovan allele density (see Fig S4). A) A visual representation of the haplotypes, with black representing an alternative allele and white representing the reference allele. Each horizontal line represents a haplotype, and is color-coded by population - the haplotypes are ordered in increasing distance from the Denisovan haplotypes. The Colombian haplotype specifically is shown in green, and seems to be most similar to other African haplotypes and not the Denisovan haplotype. B) A plot illustrating the distance of each of the haplotypes from the Denisovan reference.

**Fig S6:**
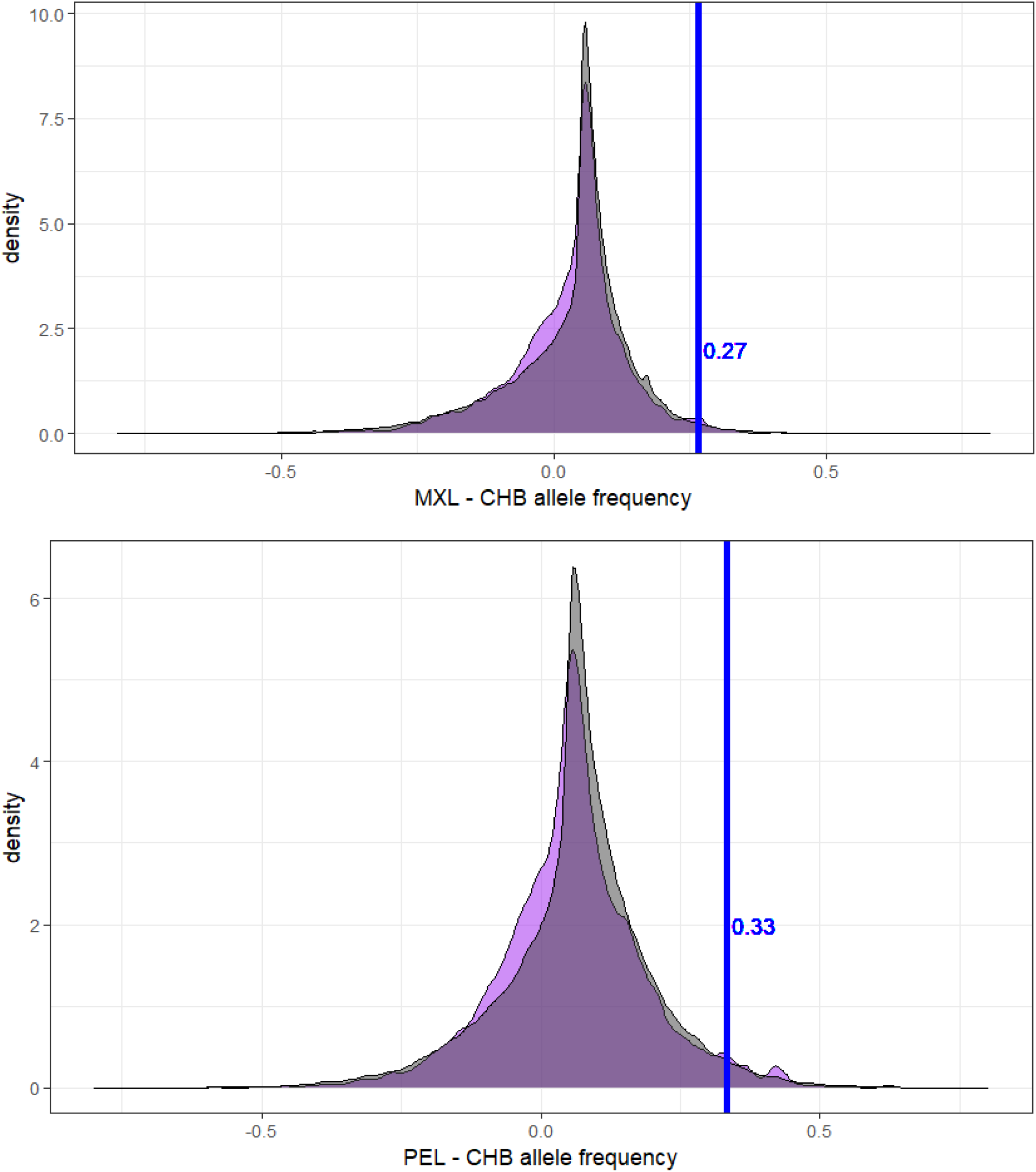
Density plots. Non-archaic rare in Africa sites in grey, archaic rare in Africa sites in purple. The top plot shows the difference in allele frequency between MXL and CHB, the bottom plot is the allele frequency difference between PEL and CHB. Both plots show only SNPs with an archaic allele frequency greater than 5%. The blue line is the value of the random sites mean plus two standard deviations.

**Table S1:**
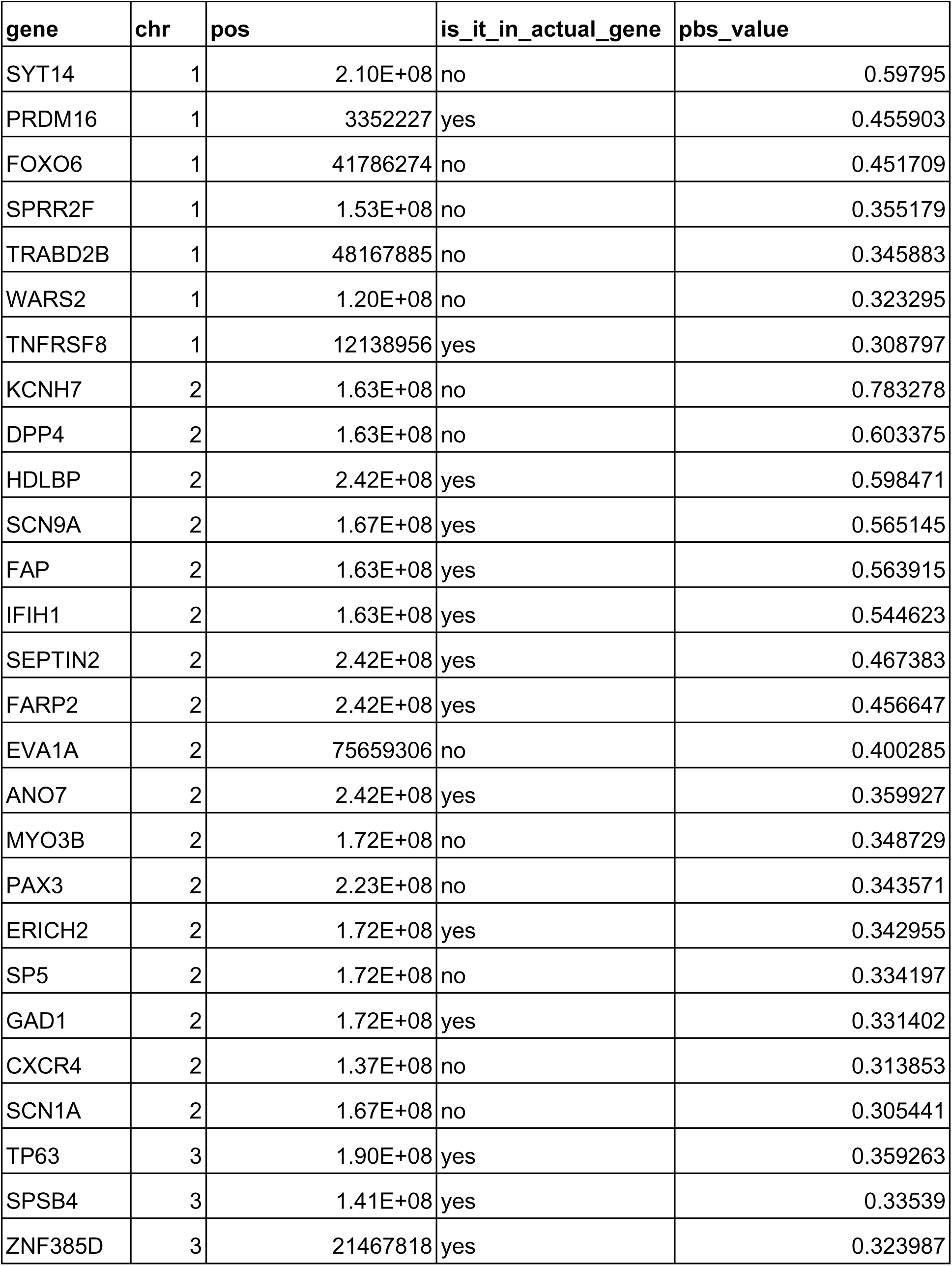
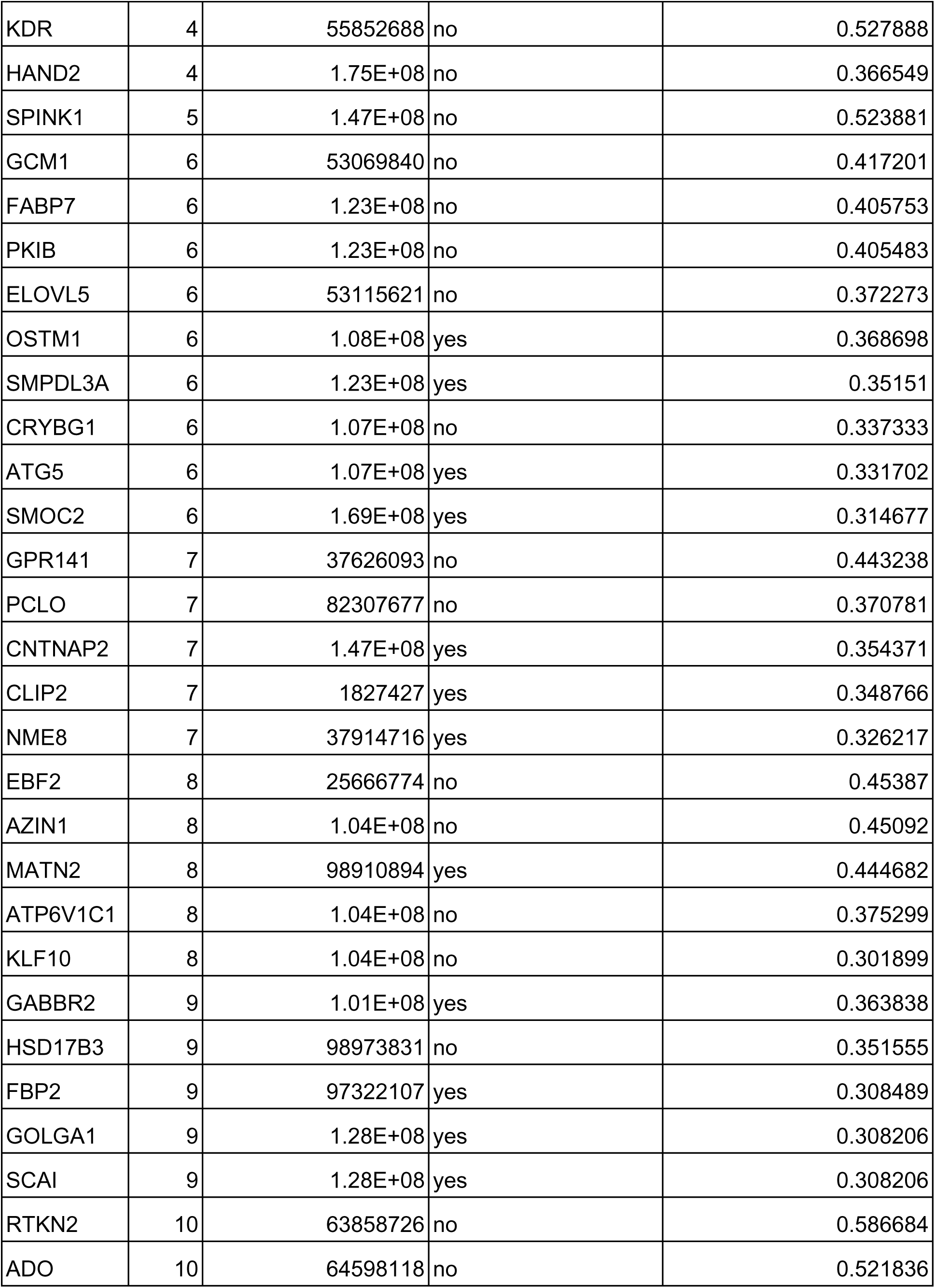
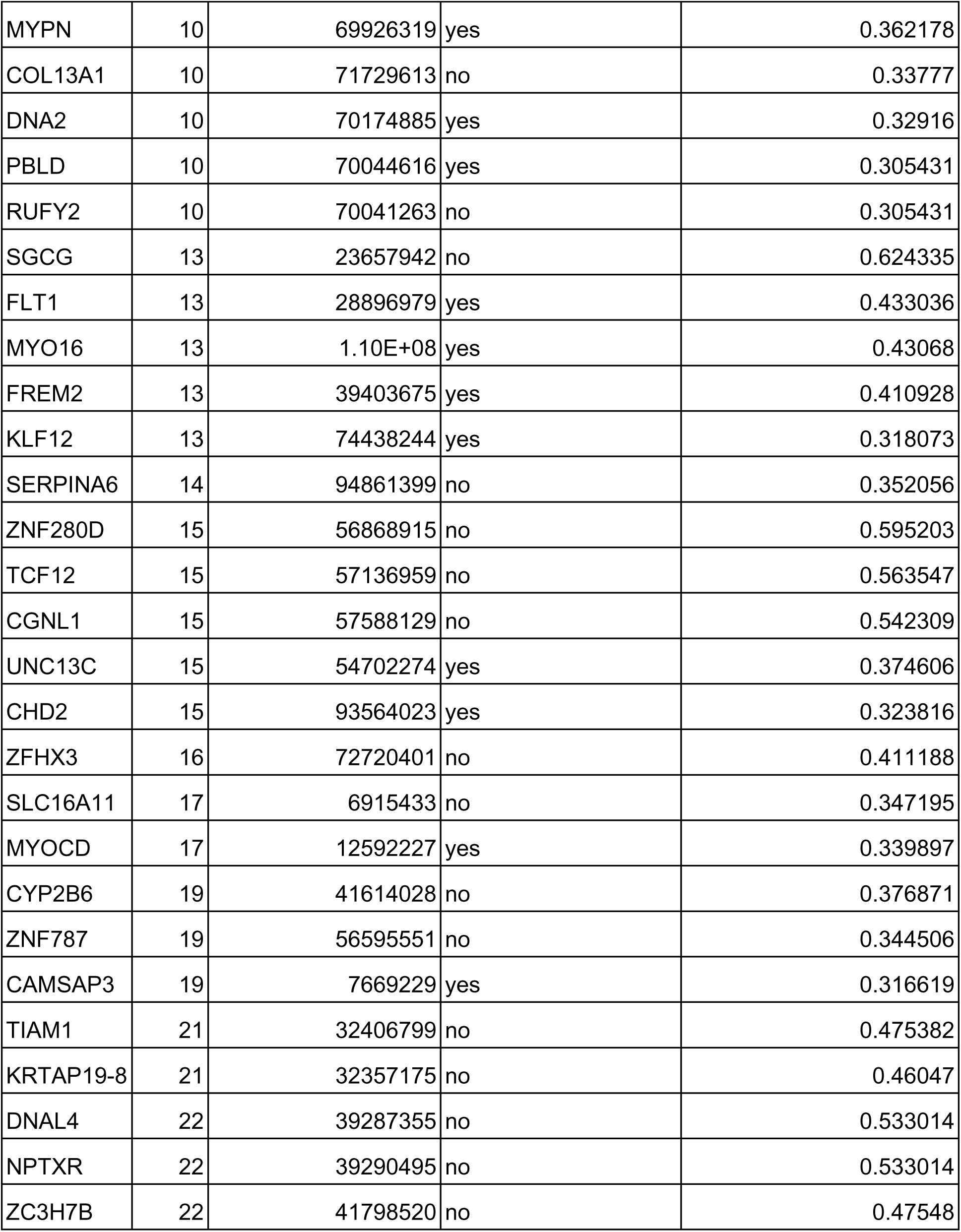
A table listing all of the SNPs with a PBS value in the top 1% of genome-wide PBS values between MXL and CHB or PEL and CHB. The closest gene is listed, along with the chromosome and position, as well as the PBS value. It is also noted whether the SNP is within the gene itself, or simply within 50 kB of that gene

## Notes

### Competing Interest Statement

The authors have declared no competing interest.

## References

1. 1000 Genomes Project Consortium, Auton A, Brooks LD, Durbin RM, Garrison EP, Kang HM, Korbel JO, Marchini JL, McCarthy S, McVean GA, et al. 2015. A global reference for human genetic variation. Nature 526:68–74.

2. Ávila-Arcos MC, McManus KF, Sandoval K, Rodríguez-Rodríguez JE, Villa-Islas V, Martin AR, Luisi P, Peñaloza-Espinosa RI, Eng C, Huntsman S, et al. 2020. Population History and Gene Divergence in Native Mexicans Inferred from 76 Human Exomes. Mol. Biol. Evol. 37:994–1006.

3. Boca SM, Huang L, Rosenberg NA. 2020. On the heterozygosity of an admixed population. J. Math. Biol. 81:1217–1250.

4. Browning SR, Browning BL, Zhou Y, Tucci S, Akey JM. 2018. Analysis of Human Sequence Data Reveals Two Pulses of Archaic Denisovan Admixture. Cell 173:1–9.

5. Bryc K, Durand EY, Macpherson JM, Reich D, Mountain JL. 2015. The genetic ancestry of African Americans, Latinos, and European Americans across the United States. Am. J. Hum. Genet. 96:37–53.

6. Bustamante CD, Henn BM. 2010. Human origins: Shadows of early migrations. Nature 468:1044–1045.

7. Coll Macià M, Skov L, Peter BM, Schierup MH. 2021. Different historical generation intervals in human populations inferred from Neanderthal fragment lengths and patterns of mutation accumulation. bioRxiv [Internet]. Available from: http://dx.doi.org/10.1101/2021.02.25.432907

8. Corbett-Detig R, Nielsen R. 2017. A Hidden Markov Model Approach for Simultaneously Estimating Local Ancestry and Admixture Time Using Next Generation Sequence Data in Samples of Arbitrary Ploidy. PLoS Genet. 13:e1006529.

9. DeGiorgio M, Jakobsson M, Rosenberg NA. 2009. Explaining worldwide patterns of human genetic variation using a coalescent-based serial founder model of migration outward from Africa. Proc. Natl. Acad. Sci. U. S. A. 106:16057–16062.

10. Dulik MC, Zhadanov SI, Osipova LP, Askapuli A, Gau L, Gokcumen O, Rubinstein S, Schurr TG. 2012. Mitochondrial DNA and Y chromosome variation provides evidence for a recent common ancestry between Native Americans and Indigenous Altaians. Am. J. Hum. Genet. 90:229–246.

11. Green RE, Krause J, Briggs AW, Maricic T, Stenzel U, Kircher M, Patterson N, Li H, Zhai W, Fritz MH-Y, et al. 2010. A Draft Sequence of the Neandertal Genome. Science 328:710– 722.

12. Han E, Carbonetto P, Curtis RE, Wang Y, Granka JM, Byrnes J, Noto K, Kermany AR, Myres NM, Barber MJ, et al. 2017. Clustering of 770,000 genomes reveals post-colonial population structure of North America. Nat. Commun. 8:14238.

13. Harris K, Nielsen R. 2016. The Genetic Cost of Neanderthal Introgression. Genetics 203:881– 891.

14. Hellenthal G, Busby GBJ, Band G, Wilson JF, Capelli C, Falush D, Myers S. 2014. A genetic atlas of human admixture history. Science 343:747–751.

15. Huerta-Sánchez E, Jin X, Asan, Bianba Z, Peter BM, Vinckenbosch N, Liang Y, Yi X, He M, Somel M, et al. 2014. Altitude adaptation in Tibetans caused by introgression of Denisovan-like DNA. Nature 512:194–197.

16. Kidd JM, Gravel S, Byrnes J, Moreno-Estrada A, Musharoff S, Bryc K, Degenhardt JD, Brisbin A, Sheth V, Chen R, et al. 2012. Population genetic inference from personal genome data: impact of ancestry and admixture on human genomic variation. Am. J. Hum. Genet. 91:660–671.

17. Korunes KL, Goldberg A. 2021. Human genetic admixture. PLoS Genet. 17:e1009374.

18. Li H, Durbin R. 2011. Inference of human population history from individual whole-genome sequences. Nature 475:493–496.

19. Mafessoni F, Grote S, de Filippo C, Slon V, Kolobova KA, Viola B, Markin SV, Chintalapati M, Peyrégne S, Skov L, et al. 2020. A high-coverage Neandertal genome from Chagyrskaya Cave. Proc. Natl. Acad. Sci. U. S. A. [Internet]. Available from: http://dx.doi.org/10.1073/pnas.2004944117

20. Malyarchuk B, Derenko M, Denisova G, Maksimov A, Wozniak M, Grzybowski T, Dambueva I, Zakharov I. 2011. Ancient links between Siberians and Native Americans revealed by subtyping the Y chromosome haplogroup Q1a. J. Hum. Genet. 56:583–588.

21. Maples BK, Gravel S, Kenny EE, Bustamante CD. 2013. RFMix: a discriminative modeling approach for rapid and robust local-ancestry inference. Am. J. Hum. Genet. 93:278–288.

22. Marnetto D, Huerta-Sánchez E. 2017. Haplostrips : revealing population structure through haplotype visualization.Price S, editor. Methods Ecol. Evol. 8:1389–1392.

23. Martin AR, Gignoux CR, Walters RK, Wojcik GL, Neale BM, Gravel S, Daly MJ, Bustamante CD, Kenny EE. 2017. Human Demographic History Impacts Genetic Risk Prediction across Diverse Populations. Am. J. Hum. Genet. 100:635–649.

24. Mautz BS, Hellwege JN, Li C, Xu Y, Zhang S, Denny JC, Roden DM, McGregor TL, Velez Edwards DR, Edwards TL. 2019. Temporal changes in genetic admixture are linked to heterozygosity and health diagnoses in humans. bioRxiv [Internet]:697581. Available from: https://www.biorxiv.org/content/10.1101/697581v1.abstract

25. Meyer M, Kircher M, Gansauge M-T, Li H, Mallick S, Schraiber JG, Jay F, Prüfer K, De C, Sudmant PH, et al. 2012. A High Coverage Genome Sequence From an Archaic Denisovan Individual. Science 338:222–226.

26. Moreno-Estrada A, Gravel S, Zakharia F, McCauley JL, Byrnes JK, Gignoux CR, Ortiz-Tello PA, Martínez RJ, Hedges DJ, Morris RW, et al. 2013. Reconstructing the population genetic history of the Caribbean. PLoS Genet. 9:e1003925.

27. Moreno-Mayar JV, Potter BA, Vinner L, Steinrücken M, Rasmussen S, Terhorst J, Kamm JA, Albrechtsen A, Malaspinas A-S, Sikora M, et al. 2018. Terminal Pleistocene Alaskan genome reveals first founding population of Native Americans. Nature 553:203–207.

28. Ongaro L, Scliar MO, Flores R, Raveane A, Marnetto D, Sarno S, Gnecchi-Ruscone GA, Alarcón-Riquelme ME, Patin E, Wangkumhang P, et al. 2019. The Genomic Impact of European Colonization of the Americas. Curr. Biol. 29:3974–3986.e4.

29. Prüfer K, Filippo C de, Grote S, Mafessoni F, Korlević P, Hajdinjak M, Vernot B, Skov L, Hsieh P, Peyrégne S, et al. 2017. A high-coverage Neandertal genome from Vindija Cave in Croatia. Science 1887:eaao1887.

30. Prüfer K, Racimo F, Patterson N, Jay F, Sankararaman S, Sawyer S, Heinze A, Renaud G, Sudmant PH, de Filippo C, et al. 2014. The complete genome sequence of a Neanderthal from the Altai Mountains. Nature 505:43–49.

31. Racimo F, Gokhman D, Fumagalli M, Ko A, Hansen T, Moltke I, Albrechtsen A, Carmel L, Huerta-s E. 2018. Archaic Adaptive Introgression in TBX15 / WARS2 Article Fast Track. 34:509–524.

32. Racimo F, Marnetto D, Huerta-Sánchez E. 2017. Signatures of archaic adaptive introgression in present-day human populations. Mol. Biol. Evol. 34:296–317.

33. Racimo F, Sankararaman S, Nielsen R, Huerta-Sánchez E. 2015. Evidence for archaic adaptive introgression in humans. Nat. Rev. Genet. 16:359–371.

34. Raghavan M, DeGiorgio M, Albrechtsen A, Moltke I, Skoglund P, Korneliussen TS, Gronnow B, Appelt M, Gullov HC, Friesen TM, et al. 2014. The genetic prehistory of the New World Arctic. Science 345:1255832–1255832.

35. Ramachandran S, Deshpande O, Roseman CC, Rosenberg N a., Feldman MW, Cavalli-Sforza LL. 2005. Support from the relationship of genetic and geographic distance in human populations for a serial founder effect originating in Africa. Proc. Natl. Acad. Sci. U. S. A. 102:15942–15947.

36. Reich D, Green RE, Kircher M, Krause J, Patterson N, Durand EY, Viola B, Briggs AW, Stenzel U, Johnson PLF, et al. 2010. Genetic history of an archaic hominin group from Denisova Cave in Siberia. Nature 468:1053–1060.

37. Reich D, Patterson N, Kircher M, Delfin F, Nandineni MR, Pugach I, Ko AMS, Ko YC, Jinam TA, Phipps ME, et al. 2011. Denisova admixture and the first modern human dispersals into Southeast Asia and Oceania. Am. J. Hum. Genet. 89:516–528.

38. Reynolds AW, Mata-Míguez J, Miró-Herrans A, Briggs-Cloud M, Sylestine A, Barajas-Olmos F, Garcia-Ortiz H, Rzhetskaya M, Orozco L, Raff JA, et al. 2019. Comparing signals of natural selection between three Indigenous North American populations. Proc. Natl. Acad. Sci. U. S. A. 116:9312–9317.

39. Rudan I. 2006. Health effects of human population isolation and admixture. Croat. Med. J. 47:526–531.

40. Sankararaman S, Mallick S, Dannemann M, Prüfer K, Kelso J, Pääbo S, Patterson N, Reich D. 2014. The genomic landscape of Neanderthal ancestry in present-day humans. Nature 507:354–357.

41. Sankararaman S, Mallick S, Patterson N, Reich D. 2016. The Combined Landscape of Denisovan and Neanderthal Ancestry in Present-Day Humans. Curr. Biol. 26:1241–1247.

42. Sankararaman S, Sridhar S, Kimmel G, Halperin E. 2008. Estimating local ancestry in admixed populations. Am. J. Hum. Genet. 82:290–303.

43. Sikora M, Pitulko VV, Sousa VC, Allentoft ME, Vinner L, Rasmussen S, Margaryan A, de Barros Damgaard P, de la Fuente C, Renaud G, et al. 2019. The population history of northeastern Siberia since the Pleistocene. Nature 570:182–188.

44. Tang H, Choudhry S, Mei R, Morgan M, Rodriguez-Cintron W, Burchard EG, Risch NJ. 2007. Recent genetic selection in the ancestral admixture of Puerto Ricans. Am. J. Hum. Genet. 81:626–633.

45. Vernot B, Akey JM. 2015. Complex history of admixture between modern humans and Neandertals. Am. J. Hum. Genet. 96:448–453.

46. Vernot B, Tucci S, Kelso J, Schraiber JG, Wolf AB, Gittelman RM, Dannemann M, Grote S, McCoy RC, Norton H, et al. 2016. Excavating Neandertal and Denisovan DNA from the genomes of Melanesian individuals. Science 352:235–239.

47. Wall JD, Yang M a., Jay F, Kim SK, Durand EY, Stevison LS, Gignoux C, Woerner A, Hammer MF, Slatkin M. 2013. Higher levels of neanderthal ancestry in East asians than in europeans. Genetics 194:199–209.

48. Wang K, Mathieson I, O’Connell J, Schiffels S. 2020. Tracking human population structure through time from whole genome sequences. PLoS Genet. 16:e1008552.

49. Witt KE, Villanea F, Loughran E, Zhang X, Huerta-Sanchez E. 2022. Apportioning archaic variants among modern populations. Philos. Trans. R. Soc. Lond. B Biol. Sci. 377:20200411.

50. Yi X, Liang Y, Huerta-Sanchez E, Jin X, Cuo ZXP, Pool JE, Xu X, Jiang H, Vinckenbosch N, Korneliussen TS, et al. 2010. Sequencing of 50 human exomes reveals adaptation to high altitude. Science 329:75–78.

51. Zhang X, Kim B, Lohmueller KE, Huerta-Sánchez E. 2020. The Impact of Recessive Deleterious Variation on Signals of Adaptive Introgression in Human Populations. Genetics 215:799– 812.

52. Zhang X, Witt KE, Bañuelos MM, Ko A, Yuan K, Xu S, Nielsen R, Huerta-Sanchez E. 2021. The history and evolution of the Denisovan-EPAS1 haplotype in Tibetans. Proceedings of the National Academy of Sciences 118:e2020803118.

